# Pre-anaesthetic anxiety phenotype stratifies anaesthesia EEG, acquired navigation, and post-anaesthesia sleep in sevoflurane-exposed mice

**DOI:** 10.64898/2026.07.13.738262

**Authors:** A. Altunkaya, M. Vadkertiova, A. Kreitmaier, C. Hedenig, S. Wipper, M.V. Schmidt, M. Kreuzer, G. Rammes, G. Schneider, T. Fenzl

## Abstract

Preoperative anxiety is common and has been associated with postoperative delirium, yet whether an anxious phenotype changes the brain’s response to anaesthesia is unknown. Here, 42 mice are classified as low-anxiety (LA) or high-anxiety (HA) by fear-retrieval freezing and exposed to sevoflurane without surgery. When adequate anaesthetic effects are reached, HA mice show a higher aperiodic spectral edge frequency than LA mice. After anaesthesia, both groups retain choice accuracy in the Water Cross Maze, but in the same trials decomposed into behavioural syllables, the learned navigation sequence is selectively disrupted in HA mice. Rapid eye-movement sleep decreases in HA mice and increases in LA mice. A single anxiety phenotype defined before induction therefore tracks three responses to the same sevoflurane exposure. These findings suggest that pre-anaesthetic anxiety marks individual vulnerability to anaesthesia in mice. The results may pave the way to interventions preserving cognitive function and sleep during perioperative recovery.

## Introduction

Anxiety is a common response to perceived threat and uncertainty, and among the most prevalent conditions in clinical settings ^1, 2^. Preoperative anxiety (POA) affects up to 80% of patients, depending on the assessment method and population studied ^3, 4^. More than 300 million surgical procedures are performed worldwide each year ^5^. Perioperative neurocognitive disorders (PND) span acute delirium to persistent cognitive decline and are a significant burden on patient outcomes and healthcare systems ^6^. Among the perioperative risk factors identified to date, POA is of particular interest because it is both prevalent and, in principle, identifiable before the procedure ^7, 8^. Several studies have linked POA to increased risk of postoperative delirium ^3, 9^. However, the relationship remains difficult to establish in clinical populations, where assessments rely on subjective self-report tools and studies are confounded by heterogeneous surgical procedures, anaesthetic protocols, and patient demographics ^10, 11^.

Animal models can address these limitations by providing a controlled environment for investigating the relationship between anxiety and anaesthetic outcome, even though they cannot recreate the full clinical picture of PND in humans ^12, 13^. In rodent models of postoperative neurocognitive disorder, a surgical trauma is added to the anaesthetic, so the effect of the anaesthetic is measured together with the inflammatory and neuroendocrine response to tissue injury ^12, 14, 15^. In adult and aged rodents, a volatile anaesthetic given on its own has altered spatial memory in some studies and left memory unchanged in others ^16, 17^, and has altered sleep/wake architecture ^18^. When anaesthetic exposure is isolated from surgery, different drugs produce distinct cognitive recovery profiles ^19^. However, anaesthetic outcomes also vary considerably between individuals ^20, 21^, and pre-existing biological vulnerability is likely to contribute to this variability. In both kinds of study the animals have been analysed as one group, and this interindividual susceptibility has not yet been incorporated into animal models of anaesthetic outcome ^13, 22^. Anxiety is one such vulnerability: it is highly prevalent in surgical populations ^3^ and a candidate moderator of anaesthetic outcome, yet it has not been tested as such in controlled animal models.

In rodents, anxiety-related behaviour is inferred from defensive responses to perceived threat and can be quantified through standardized behavioural paradigms ^23, 24^. Individual differences in this dimension have been studied extensively through selectively bred high-anxiety and low-anxiety lines ^25, 26^. However, these lines represent artificial extremes of the anxiety spectrum and may not capture the range of anxiety variation present in genetically diverse populations ^27, 28^. Data-driven classification methods can resolve anxiety-related phenotypes within standard inbred strains from behavioural data alone ^27, 29^. It remains untested whether these phenotypes relate to differential vulnerability beyond the anxiety domain, including the response to general anaesthesia.

Sevoflurane is the most widely used volatile anaesthetic in current clinical practice ^30^. In rodent models, sevoflurane produces dose-dependent changes in cortical activity. At surgical tolerance, the electroencephalogram (EEG) shows burst suppression, a pattern in which high-voltage bursts alternate with near-silent periods ^31^. Beyond the intra-anaesthetic period, exposure to volatile anaesthetics has been associated with post-anaesthetic cognitive effects in rodent models ^32, 33^. Volatile anaesthetic exposure has also been shown to alter sleep/wake architecture, with rebound of non-rapid eye movement sleep (NREMS) ^18^ and rapid eye movement sleep (REMS) reported in rodents following sevoflurane, isoflurane, and halothane exposure ^34, 35^. Intra-anaesthetic brain activity, post-anaesthetic spatial cognition, and post-anaesthetic sleep together define three outcome domains for assessing the effects of general anaesthesia, yet whether pre-existing anxiety phenotype modulates any of these outcomes has not been tested.

In rodent behavioural neuroscience, complex behavioural sequences are routinely reduced to summary metrics such as latencies, distances, and error counts ^36^. These measures capture task-level performance but collapse the temporal and sequential structure of behaviour into single values ^37, 38, 39^. Unsupervised behavioural decomposition can instead segment continuous movement into recurring behavioural syllables without predefined categories, capturing both the repertoire and the sequential organization of behaviour ^40, 41^. Whether such an approach can uncover individual differences in the behavioural response to general anaesthesia has not been explored.

Here, we classified 42 C57BL/6N mice as low-anxiety or high-anxiety by unsupervised clustering on fear retrieval behaviour and examined intra-anaesthetic EEG activity, spatial cognition before and after sevoflurane anaesthesia, and sleep/wake architecture across baseline and post-anaesthetic recordings. We asked whether a pre-anaesthetic anxiety phenotype shapes the response to sevoflurane across all three domains: whether high-anxiety and low-anxiety mice differ in the cortical EEG at anaesthetic effect, whether their learned navigation behaviour separates after anaesthesia, and whether their post-anaesthetic sleep/wake responses diverge by group.

## Methods

### Animals

42 male C57BL/6N mice (Charles River Laboratories, Sulzfeld, Germany), aged 12-16 weeks, were used. The care of laboratory animals and all experiments were performed in accordance with the recommendations of the European Union and the ARRIVE guidelines 2.0 ^42^. All experimental protocols received authorization from the Bavarian Government (ROB-55.2-2532.Vet_02-21-73). A minimum sample size of 16 mice per group was set by a power calculation in the licence application for a two-group comparison with rapid eye movement sleep as the primary outcome (significance level 0.05, power 0.80, Cohen’s d = 1.1). Mice were housed individually in recording cages within sound-attenuated chambers on an inverted 12 h light-dark cycle (lights off at 08:00, Zeitgeber time ZT0; lights on at 20:00, ZT12) at 22 ± 2°C and 55 ± 10% humidity. Food and water were provided ad libitum. Specific details of the housing, enrichment, and husbandry protocols were identical to those described in previous work from our laboratory ^18, 43^. The mice were run in 5 sequential batches (1 × 10, 4 × 8 mice).

### Experimental timeline

Each mouse underwent a 23-day protocol spanning surgery, recovery, behavioural phenotyping, spatial learning, anaesthesia, and post-anaesthetic assessment (Fig. 1a). Stereotactic electrode implantation was performed on D (Day) −15, followed by a 14-day recovery period (D−14 to D0). Baseline EEG/electromyography (EMG) was recorded for 24 hours on D1 (bEEG). Fear conditioning (FC) was performed on D2 and fear retrieval (RET) on D4, with a rest day between. Water Cross Maze (WCM) training spanned D7-D11 (5 consecutive days), followed by 2 rest days and a pre-anaesthesia Test session on D14 (10 trials). Sevoflurane anaesthesia was administered on D16. Post-anaesthesia ReTest was performed on D17 (10 trials), and 24-hour post-general anaesthesia (GA) EEG was recorded on D18. All behavioural experiments (FC, RET, WCM) were conducted at ZT14 (6 hours into the dark/active phase) in an isolated testing room. Experimental anaesthesia began at ZT13 for the first mouse, with subsequent mice at ZT14.5, ZT16, and ZT17.5 (4 mice per day, consecutive). The testing order of the mice was not randomized.

**Figure 1:**
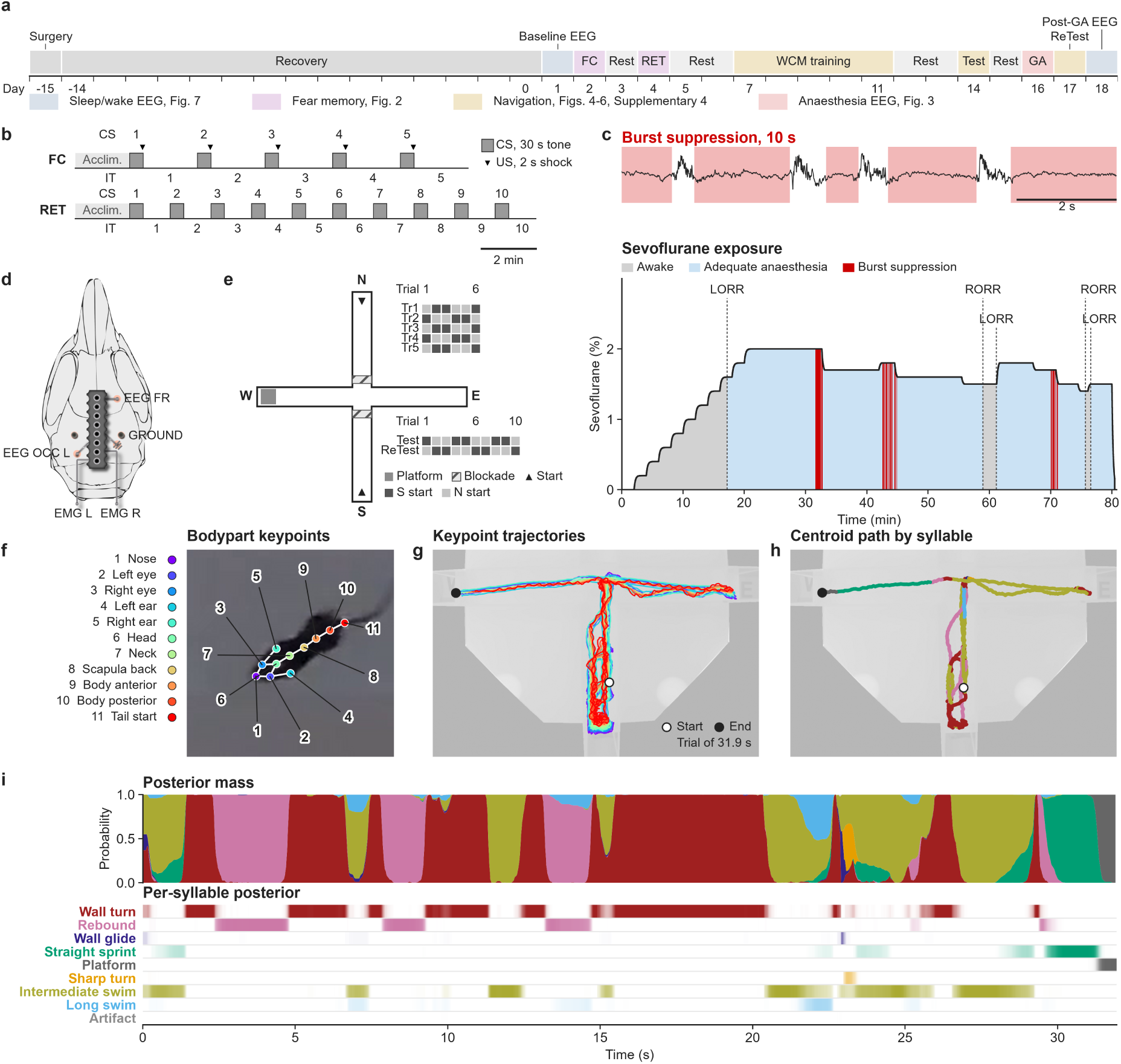
Experimental design, recordings and the behavioural syllable pipeline. (a) Experimental timeline from electrode implantation (day −15) to the 24-hour post-anaesthesia EEG (day 18): baseline EEG, fear conditioning (FC), fear retrieval (RET), Water Cross Maze (WCM) training, Test, sevoflurane general anaesthesia (GA) and ReTest; colours mark the figures in which each phase is analysed. (b) FC and RET protocols: a 2 min acclimation period, then five (FC) or ten (RET) 30 s conditioned stimuli (CS), each followed by a 2 min (FC) or 1 min (RET) intertone interval (IT); in FC the unconditioned stimulus (US, 2 s foot shock) co-terminates with each CS. (c) Sevoflurane concentration over one experimental anaesthesia session, shaded by state: awake, Adequate anaesthesia between loss of righting reflex (LORR) and return of righting reflex (RORR), and burst suppression; Inset: 10 s of EEG during burst suppression, suppression periods shaded. (d) EEG/EMG implant on the dorsal skull: epidural EEG electrodes over frontal (EEG FR) and occipital (EEG OCC L) cortex, contralateral ground, nuchal EMG electrodes (EMG L, EMG R) and the 8-pin socket. Mouse skull adapted from Cook ^44^. (e) Water Cross Maze with the hidden platform in the W arm and the blockade positions; tiles give the planned start arm (N or S) of every trial on training days Tr1-Tr5 (6 trials per day), Test and ReTest (10 trials each). (f) The 11 DeepLabCut bodyparts, numbered, on one video frame. (g) Trajectories of the 11 bodyparts over one 31.9 s trial, from start (open circle) to platform arrival (filled circle). (h) Centroid path of the same trial coloured by the most probable Keypoint-MoSeq syllable of each frame, from the marginal probabilities in (i). (i) Gibbs marginal probabilities of the 8 merged syllables and the artifact category over the same trial, as stacked posterior mass and as per-syllable posterior.

### EEG/EMG Implantation and Recording

Under isoflurane anaesthesia, mice were implanted with chronic EEG/EMG electrodes using a stereotaxic frame (Leica Mikrosysteme Vertrieb GmbH, Wetzlar, Germany). Two epidural EEG gold wire electrodes (150 μm diameter, 751 GG; C. Hafner, Pforzheim, Germany; Fig. 1d) were placed over frontal cortex (ML +1.0 mm, AP +2.8 mm, DV −0.5 mm) and occipital cortex (ML +2.8 mm, AP −3.5 mm, DV −0.5 mm). A ground electrode of the same material was placed contralaterally (ML −2.8 mm, AP −3.5 mm, DV −0.5 mm). Two gold wire EMG electrodes were inserted into the nuchal musculature for muscle tone assessment. Two stainless steel screws provided mechanical stability. All electrodes were connected to a custom 8-pin PCB socket (PRECI-DIP SA Series 861, Delémont, Switzerland) and secured with dental cement. Surgical procedures and the chronic EEG/EMG recording setup have been described previously ^43, 45, 46^.

The PCB socket was connected to a recording system consisting of a commutator and a rotatable, weight-balanced swivel system, as previously described ^43^. The raw EEG signal was amplified (INA128 instrumentation amplifier combined with an OPA244 non-inverting stage, gain ≈10,000×) and analogue band-pass filtered from 0.16 Hz to 1 kHz by a custom front end, using two cascaded single-pole high-pass sections and a 4th-order Butterworth low-pass. The EMG channel used the same architecture and the same filter corners, with a gain of 1,000×. Both signals were digitized with 16-bit resolution at a sampling rate of 256 Hz (NI USB-6343; National Instruments, Austin, TX, USA) and recorded continuously using EGErA recording software (SEGA, Cologne, Germany). Three recording sessions were used for this study: baseline sleep (bEEG, 24 h on D1), intra-anaesthetic EEG (GA-EEG, D16), and post-anaesthetic sleep (pGA-EEG, 24 h on D18).

### Fear Conditioning and Retrieval

Cued fear conditioning (FC) and fear retrieval (RET) were employed to assess and classify intrinsic anxiety phenotypes. Experiments were conducted in an isolated room away from the housing area using a fear conditioning apparatus (FREEZING AND STARTLE Threshold Sensor including Sound Attenuating Box, LE116 76-0280; Panlab, Barcelona, Spain). The floor of the conditioning box consisted of a metal grid for electric foot-shock administration, and the cage top featured a circular opening for video camera mounting, confined within the sound-attenuated box. The setup was controlled using Packwin software (Panlab, Barcelona, Spain).

The FC protocol (Fig. 1b) began with a 2-minute acclimation period, followed by a 30-second auditory conditioned stimulus (CS; 10 kHz, 75 dB). During the final 2 seconds of the CS, a co-terminating foot shock (unconditioned stimulus, US; 0.6 mA) was administered. This was followed by a 2-minute intertone interval without auditory or shock stimuli. The sequence was repeated five times, yielding 5 tone phases and 4 intertone intervals, and concluded with a 2-minute consolidation phase counted as the 5th intertone interval. The RET test was performed 48 hours post-FC at the same time and location. The protocol started with a 2-minute acclimation phase, followed by a 30-second CS (10 kHz, 75 dB) without the US and a 1-minute intertone interval. The sequence was repeated ten times, yielding 10 tone and 10 intertone intervals, and terminated with a 1-minute consolidation phase.

Freezing behaviour was scored manually from video by an experienced observer using a custom MATLAB script that recorded keyboard-triggered transitions between “freezing” and “moving” states. Freezing was defined as the absence of all movement except respiration for more than 0.5 seconds. Six freezing metrics were extracted per animal: baseline freezing (pre-tone, FC session), tone freezing (during CS, FC), intertone freezing (between CS, FC), context freezing (retrieval context, RET), pre-tone freezing (novel context before cue, RET), and cue freezing (tone in novel context, RET).

### Water Cross Maze

Spatial learning and spatial memory were assessed using a custom-built Water Cross Maze (WCM; Geißler Plexiglas, Freudenberg, Germany; Fig. 1e). The maze consisted of four arms (50 cm long, 10 cm wide) with 30 cm high walls, constructed from 0.5 cm thick clear acrylic glass ^47^. The maze was filled with fresh tap water (23°C, 11 cm depth) before each testing day. A hidden escape platform (8 × 8 cm) was submerged 1 cm below the water surface in one arm. Distal visual cues were placed around the room (the north and west arms screened with a white curtain while the east and south arms remained open to the surrounding laboratory, providing distal spatial cues, max. 14 lux at mouse level). Start arms were pseudo-randomized across trials. All trials were recorded at 1920 × 1080 resolution and 30 fps (GoPro, Inc., San Mateo, CA, USA).

Training consisted of 5 consecutive days (D7-D11) with 6 trials per day (30 trials total). Maximum trial duration was 30 seconds. If the platform was not found within this time, the mouse was guided to it by using an opaque rod held in front of the mouse, which the mouse then swam towards. Inter-trial rest was 10 minutes under a heating lamp. Each training day lasted approximately 70 minutes per mouse. Pre-anaesthesia Test (D14) and post-anaesthesia ReTest (D17) each consisted of 10 trials and lasted approximately 1 hour 45 minutes per mouse. The 3-day interval between the last training day and Test was matched to the interval between Test and ReTest (accounting for the experimental anaesthesia day). Conventional scoring metrics included escape latency, accuracy (correct/incorrect arm choice), wrong platform visits^47^ and start bias, together with two per-session classifications, accurate learner and biased starter (Supplementary Fig. 4). Start bias was the absolute difference between the number of accurate trials started from the north arm and the number started from the south arm within a session. A mouse counted as an accurate learner on a session when at least 5 of its 6 training trials, or at least 8 of its 10 Test or ReTest trials, were accurate, and as a biased starter when its start bias reached 2 on a training day or 4 at Test or ReTest.

### Experimental Anaesthesia Protocol

Sevoflurane anaesthesia was administered in a clear acrylic induction chamber (0.8 L, 160 × 120 × 135 mm; model 8329001, AgnThos, Lidingö, Sweden). The gas mixture consisted of 50% oxygen and 50% medical air (compressed, purified air; ∼21% oxygen) at a total flow rate of 4 L/min, achieving complete chamber air exchange every 12 seconds (Capnomac; Datex-Ohmeda, Helsinki, Finland). Sevoflurane (Sevorane; AbbVie, Wiesbaden, Germany) was delivered via a calibrated vaporizer (Dräger Vapour 2000; Drägerwerk AG, Lübeck, Germany).

Sevoflurane concentration (Fig. 1c) was increased stepwise in 0.2% increments every 2 minutes until reaching 2.0%. Loss of righting reflex (LORR) was assessed by gentle rotation of the chamber. The experimenter waited 30 seconds after the last visible EMG activity before tilting the chamber. LORR was defined during offline scoring of the experimental anaesthesia session. At the 20-minute mark, maintenance was started for 1 hour. During maintenance, concentration was adjusted based on live raw EEG to keep the mouse out of burst suppression: concentration was reduced if burst suppression was detected and increased if signs of lightening or movement appeared (Fig. 1c). Anaesthetic concentration was recorded at each behavioural event (LORR, burst suppression onset/offset, return of righting reflex (RORR)). At the 1-hour maintenance mark, the vaporizer was turned off. Mice remained in the chamber for 10 minutes, and all animals recovered spontaneously within this period.

Chamber temperature was maintained at 24-26°C using a heating pad beneath the box, an infrared heating lamp at a distance, and bedding material from each mouse’s own home cage placed inside the box. Temperature was monitored continuously via a thermistor placed above the bedding material.

Of the 42 mice, 34 yielded usable intra-anaesthetic EEG recordings, 22 low-anxiety (LA) and 12 high-anxiety (HA) mice. Eight mice were excluded due to a recording software failure in which live EEG was visible during the session, but the data were not saved to disk (6 HA, 2 LA). The loss was independent of group assignment. All 42 mice completed the behavioural protocols. Exclusion criteria were set before the LA and HA groups were compared. A flow diagram in the style of the Consolidated Standards of Reporting Trials detailing sample attrition across all analyses is provided in Supplementary Fig. 1. The scored anaesthetic state of every mouse across the maintenance phase is shown in Supplementary Fig. 3g.

### Sleep Scoring

EEG and EMG signals were resampled to 125 Hz and scored in 4-second epochs into three vigilance states: WAKE, non-rapid eye movement sleep (NREMS), and rapid eye movement sleep (REMS). Scoring was performed using a semi-automated vigilance scoring algorithm ^48, 49^, with manual verification and correction in the final step.

### Video Preprocessing

All WCM trial videos were processed through a two-stage pipeline. First, raw videos were compressed using FFmpeg 7.1.1 (https://ffmpeg.org) with the H.264 codec (libx264, CRF 24, preset medium, 1920 × 1080 resolution, 30 fps, audio stripped). Second, the maze region was extracted from the camera field of view using custom-written Python scripts (OpenCV) that applied automated thresholding by Otsu’s method: a median frame was computed from 50-100 frames sampled between the 10th and 90th percentile of the video, contour detection identified the maze boundary, and Hough line-based rotation correction ^50^ was applied via affine transformation. The cropped output measured 1,152 × 1,080 pixels for every video.

### DeepLabCut Pose Estimation

Markerless pose estimation was performed using DeepLabCut (DLC) 3.0.0rc13 (PyTorch backend) ^51, 52^ (Fig. 1g). A bottom-up HRNet-W32 model ^53^ was initialized from the SuperAnimal TopViewMouse backbone (DLC Model Zoo) ^54^ using transfer Mode A (backbone encoder only, random prediction head). The model was fine-tuned on 2,251 manually labelled frames from 131 videos tracking 11 bodyparts: nose, left eye, right eye, left ear, right ear, head, neck, scapula back, body anterior, body posterior, and tail start (Fig. 1f). The mid tail and the tail end were also labelled but were excluded from all downstream analysis. Training used a 95%/5% train/test split (shuffle 4), 200 epochs with batch size 24, the AdamW optimizer (learning rate 1 × 10⁻⁴, reduced to 1 × 10⁻⁵ at epoch 160 and 1 × 10⁻⁶ at epoch 190), and 16 parallel data-loading worker processes on an NVIDIA GeForce RTX 4090 GPU (NVIDIA Corporation, Santa Clara, CA, USA). The best snapshot (epoch 200) achieved a mean average precision of 97.67, a mean average recall of 98.58, and a root mean square error of 1.49 pixels on the held-out test set. A confidence threshold (p-cutoff) of 0.6 was applied for downstream filtering.

Inference was performed on all 2,092 cropped trial videos (1,152 × 1,080 pixels, 30 fps), producing per-frame x, y, and likelihood estimates for each body part in HDF5 format. Trial boundaries were then identified from these pose estimates. The start frame was defined as the first frame where at least 10 of 11 bodyparts exceeded a confidence threshold of 0.78 for at least 7 of 10 consecutive frames, offset by 3 frames. The end frame was defined by platform arrival, meaning the frame at which the mouse centroid entered the platform zone with stationarity below 15.1 pixels over 15 frames. Videos were then trimmed using FFmpeg 7.1.1, and the corresponding DeepLabCut output files in HDF5 and CSV format were sliced to matching frame ranges.

### Keypoint-MoSeq Pipeline

Behavioural syllable discovery was performed using Keypoint-MoSeq ^40, 41^ (Fig. 1h,i), which fits an autoregressive hidden Markov model to pose time series extracted by DeepLabCut. All 11 tracked bodyparts were used as input. The pipeline proceeded in three stages: principal component analysis (PCA; latent dimension = 4, whitening enabled, fitted on up to 1,000,000 frames), autoregressive HMM initialization (AR order = 3, 100 latent states, 50 iterations), and full model fitting.

A grid search was conducted across 5 AR kappa values (1 × 10⁴, 5 × 10⁴, 1 × 10⁵, 1 × 10⁶, 1 × 10⁷; full model kappa 10× lower) with 10-20 random seeds each, yielding 80 candidate models fitted for 200-300 iterations. Model selection was based on expected marginal likelihood. The selected model (AR kappa = 1 × 10⁵, full-model kappa = 1 × 10⁴, seed 3) was subjected to Gibbs sampling ^41^ (200 burn-in iterations, 100 posterior samples, 10 steps per sample, totaling 1,200 iterations) to obtain frame-level marginal probability distributions over syllable states.

The maximum a posteriori (MAP) labelling assigned at least one frame to 85 syllable states. Of these, 50 were excluded as artifacts because they occurred in fewer than 0.5% of all syllable bouts and were kinematically incoherent ^41^. The remaining 35 syllables were merged into 8 behavioural categories based on hierarchical similarity (dendrogram-based) and manual inspection using a custom-made Syllable Merge Explorer GUI. The 8 merged syllables were: Wall Glide, Straight Sprint, Sharp Turn, Rebound, Intermediate Swim, Long Swim, Platform, and Wall Turn. One additional category (ARTIFACT) included all excluded syllables.

A hard label assigns each frame to a single syllable. Gibbs marginal probabilities were used instead of MAP hard labels, so each frame held a probability distribution across all syllable states. Hard assignment was applied at exactly one point: at each frame the argmax of the 8-dimensional merged Gibbs marginal probability vector (ARTIFACT excluded) was taken, and consecutive frames sharing the same label were grouped into bouts. Each bout’s full probability vector was retained, and all downstream analyses (frequency estimation, transition probability computation, uniform manifold approximation and projection (UMAP) embedding, and statistical testing) operated on these retained soft vectors instead of the hard labels. For the transition analyses and the UMAP embedding, the Platform entry was removed from each bout vector and the remaining 7 entries were re-normalized to sum to 1. Bouts labelled Platform were left out of the UMAP embedding.

### Anxiety Phenotyping

Trait anxiety was assessed using a data-driven clustering approach on fear retrieval freezing behaviour. Four features were extracted per mouse from the retrieval session: median freezing during early tone presentations (CS1-5), late tone presentations (CS6-10), early intertone intervals (CS1-5), and late intertone intervals (CS6-10). One mouse (M4AA41) had missing retrieval data due to technical failure, and the missing values were imputed using K-nearest neighbours (K = 5, inverse distance weighting, Euclidean distance on 10 FC-derived features). Clustering was performed using k-medoids (partitioning around medoids algorithm) with Manhattan distance and k = 2. Cluster robustness was validated across 5 different algorithms: k-medoids Manhattan, k-medoids Euclidean, k-means Euclidean, Ward hierarchical, and Gaussian mixture model (full covariance). Bootstrap stability (5,000 iterations) and gap statistic (5,000 Monte Carlo references) were computed for internal validation. The stability of the group assignment of each mouse was tested in a separate bootstrap with 5,000 resamples of the 42 mice (random seed = 42). Each resample was clustered by k-medoids with Manhattan distance and k = 2, keeping the best of 20 k-medoids++ initializations. In each resample, the cluster with the higher mean freezing was labelled HA, as for the original groups. Each mouse was then assigned the label of its nearest bootstrap medoid in every resample. In a second reading, each mouse was labelled only in the resamples that included that mouse. A label permutation test (10,000 shuffles) was run on the key downstream metrics to compare the low anxiety/high anxiety (LA/HA) grouping against a null distribution. The LA and HA groups were computed after the experiments had been completed and were applied retrospectively to all analyses. Data collection and scoring were therefore blind to group.

### Intra-Anaesthetic EEG Analysis

Raw signals were preprocessed using a 3-stage asymmetric filter: (1) low-pass at 48 Hz (4th-order Butterworth, second-order sections [SOS], zero-phase via sosfiltfilt), (2) high-pass at 0.5 Hz (2nd-order Butterworth, SOS, zero-phase), and (3) 50 Hz notch filter (quality factor Q = 30, zero-phase via filtfilt). The analysis frequency range was 0.5-45 Hz.

The maintenance period was defined as the interval from the last LORR event within the first 20 minutes to the last RORR event. EEG segments were classified into four maintenance phases: Adequate, Discontinuity, Burst (burst suppression, burst epochs), and Suppression (burst suppression, suppression epochs).

Event-based metrics: Anaesthetic concentration at LORR, RORR, and four maintenance phases. Temporal metrics: Episode counts, durations, state proportions, median bout durations, and latencies for each maintenance phase. Burst suppression architecture: Suppression dominance ratio, burst suppression ratio, suppression and burst proportions, root mean square (RMS) amplitudes, and amplitude ratio. Spectral power: Welch periodogram-based absolute and relative band power in five frequency bands: delta (0.5-4 Hz), theta (4-8 Hz), alpha (8-13 Hz), beta (13-24 Hz), and gamma (24-45 Hz), computed separately for each maintenance phase.

Spectral parameterization: The fitting oscillations and one over f (FOOOF) algorithm (fooof 1.1.1) ^55^ was applied in knee aperiodic mode to decompose power spectra into aperiodic (1/f) and periodic (oscillatory peak) components. The model decomposes the power spectrum as 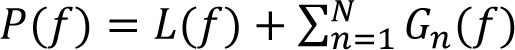. Parameters applied ^56^: peak width limits 2-12 Hz, maximum 3 peaks, minimum peak height 0.3, peak threshold 2.0, frequency range 0.5-45 Hz. Aperiodic parameters (exponent, offset, knee) and periodic parameters (peak centre frequency, power, bandwidth) were extracted per maintenance phase.

Spectral analysis on the aperiodic component: Spectral edge frequency at the 95th percentile (SEF95), defined as the frequency below which 95% of spectral power resides, 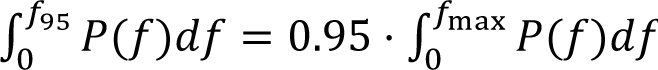 and spectral entropy were computed on the FOOOF-derived aperiodic component ^56^.

Transition dynamics: Band power, SEF95, and permutation entropy (PeEn) were computed in peri-event windows (±120 s) around LORR and RORR. The change in relative band power across LORR was the difference between the 60 s after and the 60 s before LORR. Cluster-based permutation tests (1,000 permutations, cluster-forming threshold p < 0.05) were used for statistical inference on time course data ^57^.

Entropy and complexity: Permutation entropy (embedding dimension m = 3, time delay τ = 1, normalized 0-1) and Lempel-Ziv complexity were computed on bandpass-filtered EEG for each maintenance phase.

### Baseline and Post-Anaesthetic Sleep Analysis

Sleep/wake architecture was analysed from 24-hour bEEG recordings (D1; n = 38, 22 LA, 16 HA) and 24-hour pGA-EEG recordings (D18; n = 33, 20 LA, 13 HA). Recordings were excluded where 24-hour coverage was incomplete (see Supplementary Fig. 1). EEG signals were preprocessed using the same 3-stage asymmetric filter described above (low-pass 48 Hz, high-pass 0.5 Hz, 50 Hz notch) and resampled to 125 Hz. Eight analysis modules were applied to each recording: (1) Vigilance state proportions: Percentage of time spent in WAKE, NREMS, and REMS, computed separately for the light phase (ZT12-ZT24) and dark phase (ZT0-ZT12). (2) Time course analysis: Vigilance state percentages in 2-hour bins across the 24-hour recording (12 bins total). (3) Transitions: Markov transition matrices between vigilance states, *T_ij_* = *P*(*s_t_*_+1_ = *j* ∣ *s_t_* = *i*), computed per half (light/dark). (4) Bout length: Median bout duration (in seconds) for each vigilance state per half. (5) Power spectral density: Welch periodogram-based band power in delta, theta, alpha, beta, and gamma bands during NREMS and REMS. WAKE spectra were expressed as relative power, each frequency bin as a percentage of the total power across the full frequency range. (6) Spectral parameterization (FOOOF, specparam 2.0.0rc6): Aperiodic and periodic decomposition of NREMS and REMS power spectra using the same parameters as the GA EEG analysis. (7) Sleep latency: Median of the first 5 inter-bout gaps as a proxy for sleep onset latency, per half. (8) Permutation entropy: PeEn (m = 3, τ = 1) computed per vigilance state per half.

Within-subject change analysis: For the mice that yielded usable recordings both at baseline and after anaesthesia (n = 31, 19 LA, 12 HA), within-subject change scores (Δ = pGA − bEEG) were computed for each metric. Group differences in these change scores were tested between LA and HA.

### Sleep Spindle Detection

Sleep spindles were detected using an automated algorithm ^58^ implemented in MATLAB R2024b (MathWorks, Natick, MA, USA). EEG signals during NREMS epochs were bandpass-filtered at 10-15 Hz (infinite impulse response Butterworth, stopbands at 3 Hz and 22 Hz, zero-phase via filtfilt). The RMS envelope was computed with a window of ¾ × sampling rate samples and smoothed using an exponential weighted moving average (α = 0.05). Spindles were identified as events exceeding 2.5× the mean NREMS RMS amplitude (upper threshold) with a lower inclusion threshold of 1.0× mean NREMS RMS. Event duration was constrained to 0.5-2.0 seconds, with a minimum inter-spindle interval of 0.1 seconds. Four metrics were extracted per recording hour: spindle count, spindle density (count per hour of NREMS), median spindle duration, and median normalized amplitude.

### Behavioural Syllable Analysis

Each bout retained the mean probability vector over its frames (soft-vector handling as described under Keypoint-MoSeq Pipeline). For the syllable atlas, the medoid bout of each syllable was the bout whose 12 z-scored kinematic features lay closest to the median feature vector of that syllable, and it is drawn as pose snapshots. Bout trails on the maze were drawn from the body centroid estimated by Keypoint-MoSeq at every frame of a bout (the centroid path). The four-step escape sequence (Wall Glide, Sharp Turn, Straight Sprint, Platform) is a descriptive schematic of the chain of syllables that takes a well-trained mouse from the start arm to the platform, defined from observation of trained mice, and no statistic was computed on it.

Syllable frequencies were estimated from Gibbs probabilities using fractional counting, *f_k_* = 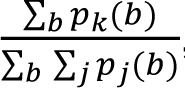, so each bout contributed to every syllable category in proportion to its probability vector.

Transition matrices were computed as soft outer products at consecutive bout boundaries, 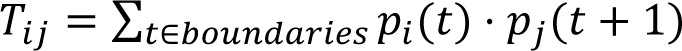 within each trial. No cross-trial concatenation was performed. Matrices were row-normalized to yield conditional transition probabilities. Comparisons between LA and HA mice were made for each cell of the transition matrix on the last training day (Tr5), at Test and at ReTest, and for within-subject change scores from Tr5 to Test and from Test to ReTest. Each change score was the transition probability on the later day minus the transition probability on the earlier day in the same mouse. In each mouse, the change from Tr5 to Test was also subtracted from the change from Test to ReTest. This difference between the two change scores was compared between LA and HA mice.

Shannon entropy of the syllable frequency vector (H = −Σ pᵢ log₂ pᵢ, in bits) was computed per mouse per day to quantify repertoire diversity. Entropy rate (H_rate_ = Σ_s_ π(s) · H(row_s_), where π(s) is the empirical frequency and H(row_s_) is the Shannon entropy of the transition row for syllable s) was computed to quantify sequential predictability. Shannon entropy and entropy rate were compared between LA and HA mice on each of the five training days (Tr1 to Tr5), at Test and at ReTest. The difference between the LA and HA syllable profiles was quantified per session as the Jensen-Shannon distance (base 2, the square root of the Jensen-Shannon divergence) between the per-syllable group medians of the per-mouse 7-syllable frequency vectors. Before the medians were taken, Platform was removed from each mouse’s frequency vector and the remaining 7 entries were re-normalized to sum to 1. For visualization of bout-level Gibbs probability structure, UMAP ^59^ was applied to the 7-dimensional merged probability vectors using precomputed Jensen-Shannon distance matrices (15 neighbours, minimum distance 0.2, random seed 42). As a linear baseline, centred log-ratio PCA ^60^ was applied to the same probability vectors.

Kinematic validation used 15 DLC-derived movement features (mean speed, bout duration, path length, displacement, sinuosity, mean elongation, mean curvature, mean head-body angle, mean angular velocity, mean acceleration, mean ear spread, and the standard deviation of speed, of curvature, of elongation and of the change in elongation between consecutive frames) normalized with a quantile transformer (uniform output distribution, random seed 42). UMAP was applied with Euclidean distance (15 neighbours, minimum distance 0.2).

### Use of Large Language Models

Large language model tools from Anthropic (Claude) and OpenAI (ChatGPT, Codex) were used for code review assistance in the analysis and figure generation scripts. The authors reviewed and verified all code and take full responsibility for the content.

### Statistical Analysis

All statistical analyses were performed in Python 3.13.0 using scipy 1.17.1, statsmodels 0.14.6, scikit-learn 1.8.0, numpy 2.4.3, and pandas 3.0.1. Group differences between LA and HA were assessed using a two-sided Mann-Whitney U test (MWU) (24 LA / 18 HA; GA subset 22 LA / 12 HA). Effect sizes: The area under the receiver operating characteristic curve (AUC) was computed as a non-parametric effect size (AUC = U / [n₁ × n₂]) with 95% confidence intervals from 10,000 bootstrap resamples (random seed = 42, percentile method).

#### Batch correction

To assess potential confounding by batch (5 sequential batches), linear mixed models (LMM) were fitted as sensitivity analyses with the formula metric ∼ group + (1|batch) using restricted maximum likelihood via statsmodels.MixedLM with L-BFGS-B optimization. The intraclass correlation coefficient 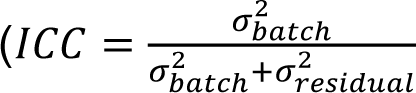 quantified the proportion of variance attributable to batch. LMM results were interpreted unidirectionally: a high ICC with non-significant LMM p-value could downgrade a MWU-significant finding, but a significant LMM could not upgrade a MWU-borderline finding.

Multiple comparison correction: The Benjamini-Hochberg false discovery rate procedure was applied within each analysis family (e.g., per GA task, per sleep analysis module) at α = 0.05. For Keypoint-MoSeq transition matrix analyses, the 49 per-cell comparisons (7 × 7 matrix) formed one correction family for each experimental day, for each of the two change scores and for the difference between the two change scores. The correction was also applied across the four transitions with an uncorrected MWU p below 0.05 at ReTest, as a separate family of four cells for each experimental day, each change score and the difference between the two change scores. Time course statistics: For peri-event EEG data (SEF95 and PeEn around LORR/RORR), cluster-based permutation tests were used to control for multiple comparisons across time points (1,000 permutations, cluster-forming threshold p < 0.05, test statistic: sum of |U − expected U| within cluster) ^57^.

## Results

### Anxiety phenotyping from fear retrieval

To define an anxiety phenotype from behaviour alone, each of the 42 C57BL/6N mice was assigned a low-anxiety (LA) or high-anxiety (HA) label from its own cued fear-retrieval performance. The resulting 24 / 18 partition was stable across methods and applied through every downstream analysis.

K-medoids clustering on the four retrieval-session freezing features separated the cohort into 24 LA and 18 HA mice (Fig. 2a). The two clusters differed in their freezing during retrieval, and HA mice already froze more than LA mice during conditioning, at all five tones and from the third intertone interval onward, with overlapping interquartile ranges (Fig. 2c,e). HA mice maintained high freezing across the whole session, both during the tones and intertone intervals, whereas LA mice spent progressively longer mobile during the intertone windows, most clearly in the late phase of the test (Fig. 2d,f). The two clusters were moderately well separated (median silhouette coefficient 0.433) (Fig. 2b), and a gap-statistic sweep revealed two clusters over the single-population null (gap = 0.256). Under bootstrap resampling of the 42 mice (5,000 resamples), 38 mice stayed in their own group in at least 69% of resamples, and four HA mice sat at the boundary between the groups (HA in 46% to 53% of resamples). Three of four alternative clustering algorithms reproduced the same 24 / 18 split, with a Gaussian mixture model giving a near-identical 22 / 20 (Supplementary Fig. 2). Fear-retrieval freezing therefore defined two stable behavioural phenotypes in a genetically uniform cohort before any anaesthetic was given.

**Figure 2:**
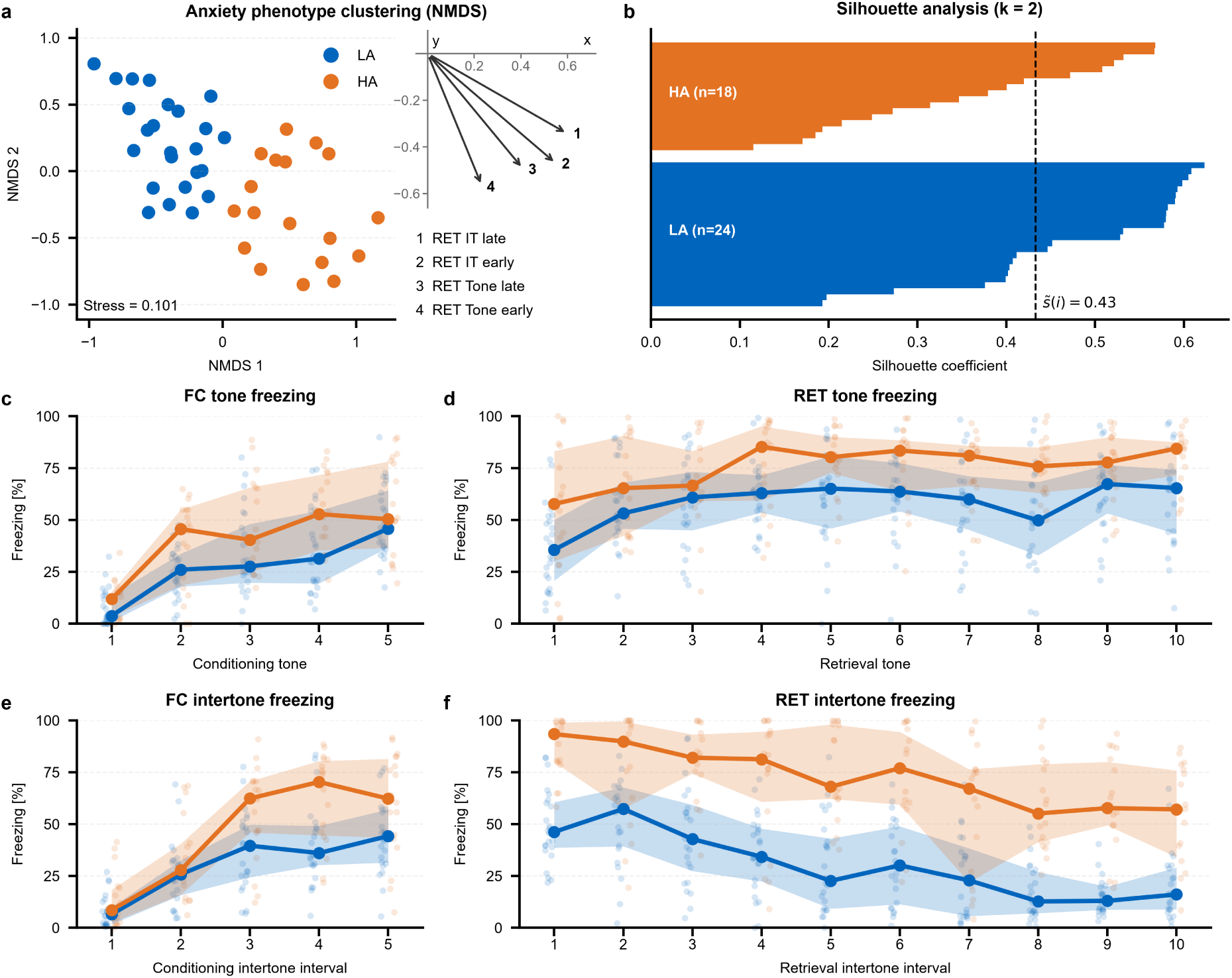
Fear-retrieval freezing separates the 42 mice into two stable anxiety phenotypes. (a) Non-metric multidimensional scaling (NMDS) of the 42 mice on Manhattan distances between the four retrieval freezing features (median freezing during CS1-5, CS6-10, ITI1-5 and ITI6-10), coloured by the k-medoids clustering (k = 2): low anxiety (LA, blue, n = 24) and high anxiety (HA, orange, n = 18). Normalised stress = 0.101. Inset: the four features as vectors in the ordination space, numbered 1-4 (1, late intertone; 2, early intertone; 3, late tone; 4, early tone). (b) Silhouette value of each mouse under the k = 2 clustering, sorted within group; dashed line, median across all 42 mice (0.43). (c-f) Freezing proportion during fear conditioning (FC; (c) five tone presentations CS1-5; (e) five intertone intervals ITI1-5) and during fear retrieval 48 h later (RET; (d) ten tone presentations CS1-10; (f) ten intertone intervals ITI1-10). Faint points, individual mice; filled circles joined by lines, group medians; shaded bands, interquartile range (25th-75th percentile) across mice.

### Intra-anaesthetic EEG signature

The two phenotypes were first compared on their cortical EEG under sevoflurane. All 42 mice received sevoflurane, and eight were excluded from the intra-anaesthetic analysis (Methods), leaving 34 (22 LA, 12 HA) in which the macro-architecture of the exposure was matched: anaesthetic concentration at loss of righting reflex (LORR) and recovery of righting reflex (RORR), time in each maintenance phase, and burst-suppression architecture did not differ between LA and HA mice (all p > 0.11; Fig. 3i, Supplementary Fig. 3a). No band-power difference reached significance in any frequency band or maintenance phase (Supplementary Fig. 3b). The largest effect was Adequate-phase absolute theta, higher in LA mice, which fell just short of significance (Mann-Whitney U test (MWU) p = 0.054). Representative frontal EEG traces distinguished the anaesthetic states (Fig. 3a). Group comparisons throughout the Results are collected in Supplementary Table 1, which reports for each measure the number of mice, the effect size with its bootstrap confidence interval, and the full set of test statistics by test.

**Figure 3:**
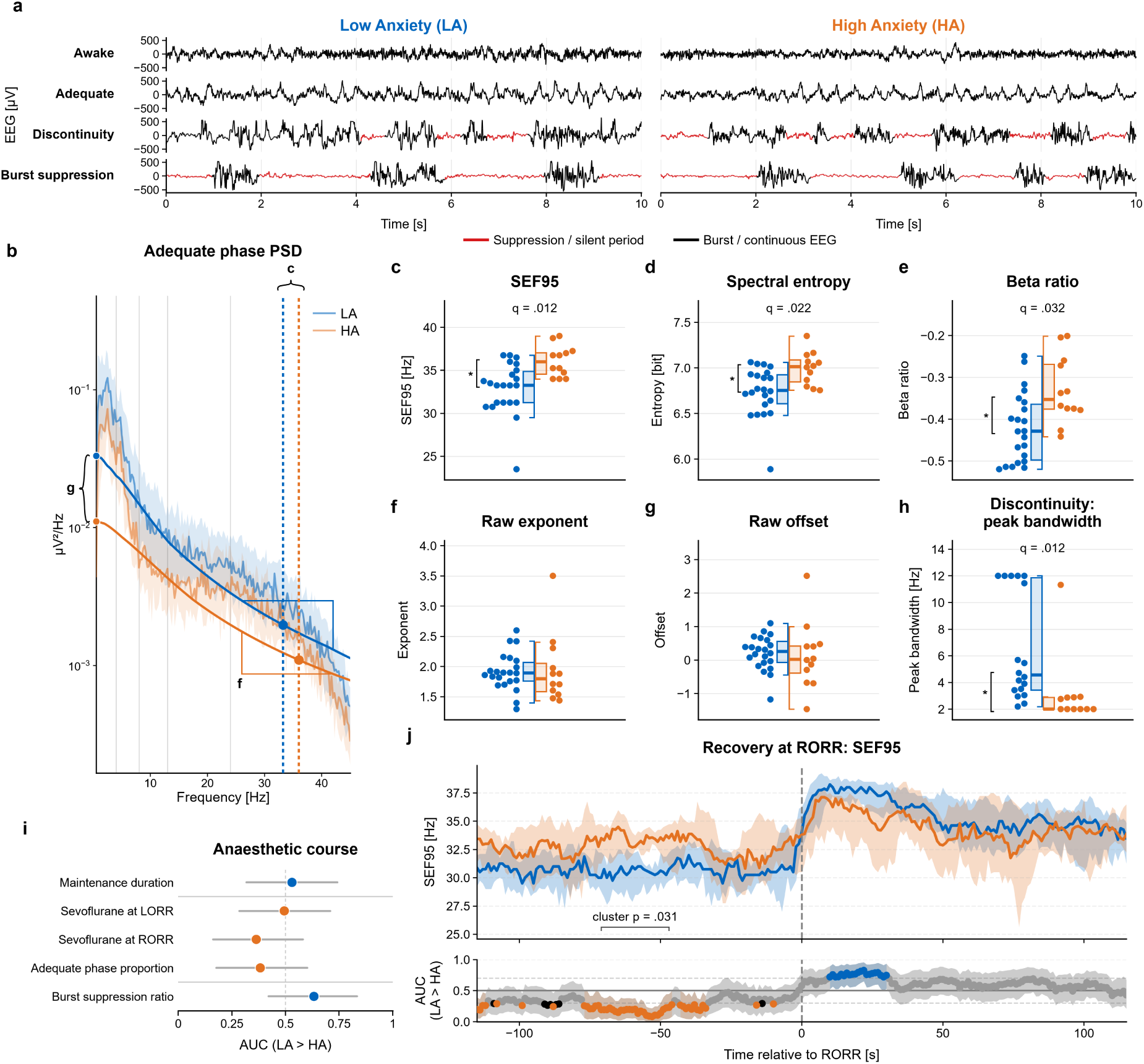
HA mice show a higher aperiodic spectral edge frequency (SEF95) than LA mice at Adequate sevoflurane depth. (a) Representative 10-s EEG segments in the four scored states (awake, Adequate anaesthesia, Discontinuity, burst suppression) for the LA and HA groups; red, suppression or silent periods; black, burst or continuous EEG. (b) Power spectral density during Adequate anaesthesia, 0.5-45 Hz (LA n = 22, HA n = 12): thin line, group median across mice; shaded band, interquartile range; thick line, median of the per-mouse aperiodic fits by fitting oscillations and one over f (FOOOF), knee model; grey rules at 4, 8, 13 and 24 Hz. Marks are lettered to the panels that quantify them: dashed rules and dots, SEF95 of each group (c); wedge over 26-42 Hz, aperiodic exponent (f); discs on the y axis, level of the aperiodic fit at 0.5 Hz (g). (c-h) Per-mouse values during Adequate anaesthesia (c-g) and Discontinuity (h): (c) SEF95, (d) spectral entropy, (e) beta ratio, (f) aperiodic exponent, (g) aperiodic offset, (h) peak bandwidth. Dots, individual mice (LA n = 22, HA n = 12 in c-g and LA n = 18, HA n = 11 in h); half-boxes, median and interquartile range, whiskers to the most extreme observation within 1.5 interquartile ranges; q, Benjamini-Hochberg FDR-corrected p of the two-sided Mann-Whitney U test; bracket with asterisk, q < 0.05. (i) AUC (LA > HA) with 95% bootstrap CI for five anaesthetic-course measures; blue, AUC above 0.5; orange, below 0.5; dashed line, AUC = 0.5. (j) SEF95 around return of righting reflex (LA n = 19, HA n = 8): pointwise median and 95% CI (mouse-resample bootstrap, 10,000 resamples); bracket, cluster from −71 to −47 s (cluster-based permutation test, p = 0.031). Lower strip, pointwise AUC (LA > HA) with 95% CI; blue and orange dots, AUC outside 0.3-0.7 with CI excluding 0.5 (LA > HA and HA > LA, respectively); black dots, AUC outside 0.3-0.7 with CI including 0.5; dashed lines, 0.3 and 0.7.

At LORR, the induction shift of cortical power toward low frequencies was complete in LA mice but attenuated in HA mice. Relative beta power fell from pre-to post-LORR in LA mice and rose in HA mice (median LA −2.3% vs HA +12.9%; MWU p = 0.010; Supplementary Fig. 3c). Relative delta power shifted further toward low-frequency dominance in LA than in HA (linear mixed model (LMM) p = 0.024; Supplementary Fig. 3c). Relative gamma change was also larger in LA (LMM p = 0.026; Supplementary Fig. 3c).

During Adequate maintenance anaesthesia, LA and HA separated most strongly in the aperiodic component of the EEG. The spectral edge frequency of the aperiodic spectrum (SEF95, the frequency below which 95% of the aperiodic power lies) was higher in HA mice (median 36.0 Hz vs 33.25 Hz; MWU p = 0.001; Fig. 3b,c). Spectral entropy on the aperiodic component moved in the same direction (LMM p = 0.010; Fig. 3d), as did the aperiodic beta ratio (LMM p = 0.014; Fig. 3e). The raw aperiodic exponent and offset trended toward a flatter slope in HA but did not reach significance (Fig. 3f,g). Permutation entropy (PeEn) and Lempel-Ziv complexity converged on the same direction (Supplementary Fig. 3e). PeEn had substantial batch variance and its group difference did not reach significance (LMM p = 0.051). Across independent modalities, HA mice showed more high-frequency aperiodic power and higher signal complexity at the same anaesthetic depth. During Discontinuity, the dominant spectral peak was broader in LA than in HA mice (median bandwidth 4.56 Hz vs 2.00 Hz; MWU p < 0.001; Fig. 3h). Batch effects across the three primary metrics were small (Supplementary Fig. 3f).

Approaching emergence, the SEF95 difference reappeared as a time course effect. In the minute before RORR, HA mice maintained a higher SEF95 than LA mice. Cluster-based permutation testing identified one significant window of 25 consecutive 1 s bins, between −71 and −47 s (cluster p = 0.031; 8 HA / 19 LA with available ±115 s windows; Fig. 3j). Non-significant clusters elsewhere in the pre-RORR window were directionally consistent. Post-RORR, the direction reversed (HA < LA) but did not survive correction. PeEn time courses moved in the same direction as SEF95 but formed no significant cluster in either the pre-RORR or post-RORR window (Supplementary Fig. 3d). The anxiety phenotype was therefore associated with a different cortical state at matched sevoflurane exposure, seen as more high-frequency aperiodic power at Adequate depth and a higher SEF95 in the minute before emergence.

### Conventional post-anaesthetic spatial metrics

Learned spatial behaviour after anaesthesia was examined next in the Water Cross Maze, first with the conventional scoring metrics. All mice acquired the task across 5 training days (Tr1-Tr5). Median escape latency fell from approximately 9.0 s [8.2; 10.3] on Tr1 to 4.4 s [4.3; 4.8] on Tr5, choice accuracy rose from approximately 50% to a median of 100% by Tr5, and wrong-platform visits declined from a median of 3 to 0 (Supplementary Fig. 4a,c,g,i; Supplementary Fig. 5). HA mice were transiently slower during early training (Tr1 MWU p = 0.019; Supplementary Fig. 4a), a ∼2 s offset that resolved by Tr3 and was absent by Tr5 (p = 0.147). On the pre-anaesthesia test session (D14), escape latency, choice accuracy, and start-arm bias did not differ between groups (Supplementary Fig. 4e,j). Wrong-platform visits showed a nominal excess in HA mice (MWU p = 0.018; Supplementary Fig. 4g,j) that did not survive family-wise correction, leaving a broadly matched baseline for the post-anaesthesia probe.

At the post-anaesthesia ReTest session (D17), conventional scoring showed no group difference beyond a small escape-latency offset. Escape latency differed statistically (LA median 4.0 s [3.7; 4.2], HA 4.3 s [3.9; 4.6]; MWU p = 0.033), but the absolute difference was 0.3 s, and both groups performed near the maximum of the task (Supplementary Fig. 4b,j). Choice accuracy reached the maximum score in both groups (medians 100%; Supplementary Fig. 4d). Wrong-platform visits and start-arm bias did not differ between groups at ReTest (Supplementary Fig. 4f,h,j). Full per-metric results are in Supplementary Fig. 5. By conventional scoring, both groups therefore retained the task after sevoflurane, and the 0.3 s latency difference was the only group difference by these metrics.

### Behavioural syllable vocabulary

To resolve how the mice moved through the maze beyond escape latency and arm choice, the same trial videos were decomposed into behavioural syllables with Keypoint-MoSeq. After excluding a pre-swim category and low-frequency outlier states, the remaining 35 states were consolidated into 8 behavioural categories covering 96.8% of frames (Fig. 4c, Supplementary Fig. 6a; exclusion thresholds in Methods).

**Figure 4:**
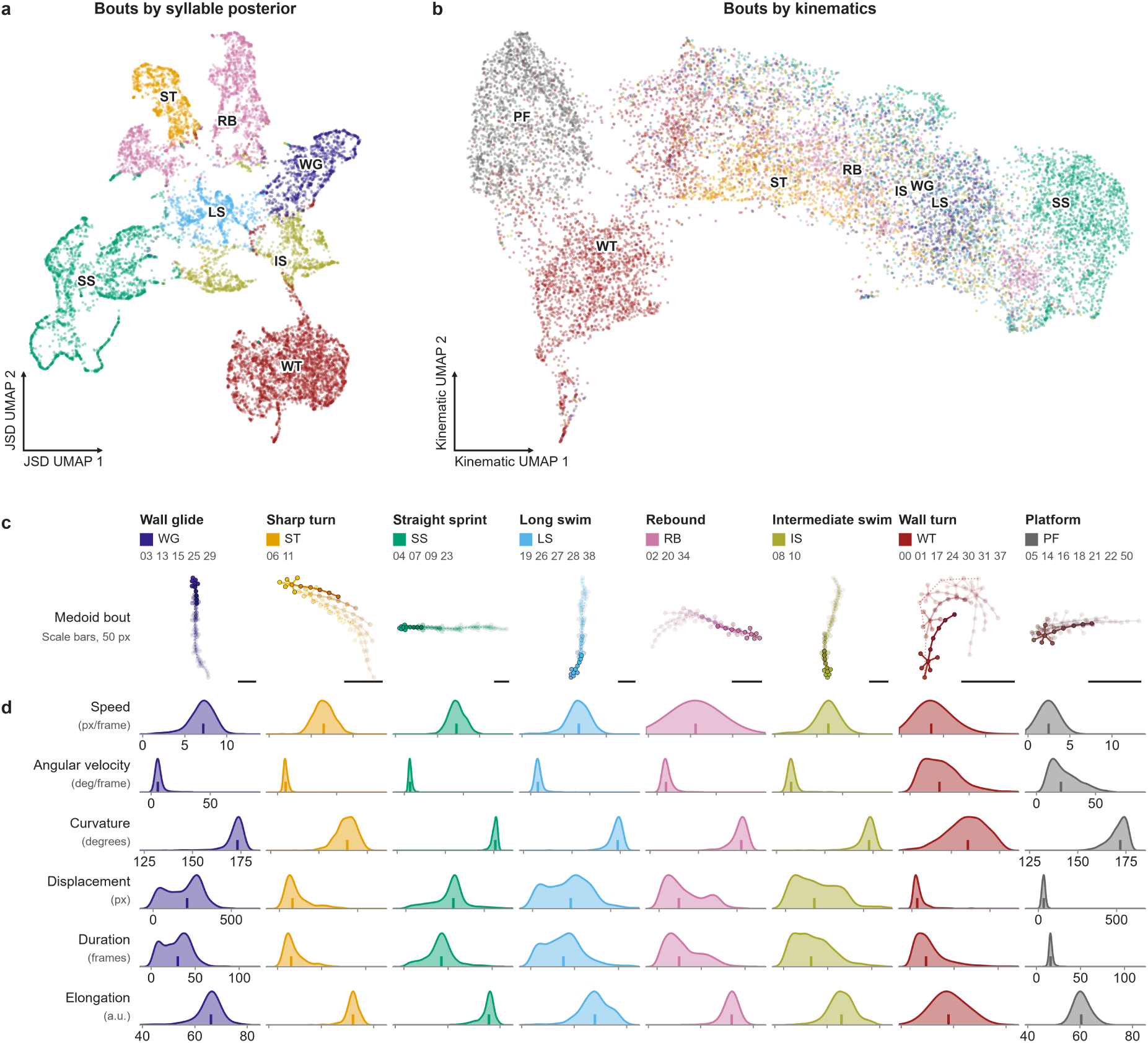
Keypoint-MoSeq resolves Water Cross Maze swimming into eight behavioural syllables. (a) Uniform manifold approximation and projection (UMAP) embedding of 12,053 bouts (42 mice, LA n = 24, HA n = 18, seven sessions) from the Jensen-Shannon distances between their merged Gibbs marginal probability vectors, coloured by syllable; Platform has no bouts in this embedding. (b) UMAP embedding (Euclidean distance) of 13,437 bouts (the same 42 mice and seven sessions) from their kinematic features, coloured by syllable. (c) The eight merged syllables: name, colour swatch with abbreviation, the syllable states merged into each (state IDs; states 12, 32, 33 and 35 are artifacts and belong to no syllable), and the medoid bout drawn as six pose snapshots fading from the first to the last frame; scale bars, 50 px. (d) Density of six kinematic features (speed, angular velocity, curvature, displacement, duration and elongation) for each syllable (the 13,437 bouts of b); vertical tick, median. Abbreviations in (a), (b) and (d) as in (c).

Intermediate Swim is an aimless search across the maze, without commitment to a specific arm. Wall Turn is a wall-bump turnaround in which the animal encounters a wall and pivots sharply. Rebound is a short post-turn acceleration that recovers swim momentum after a Sharp Turn or a Wall Turn. Long Swim is an open-arm swim of moderate duration traversing the middle of a non-goal arm without sustained wall contact. Wall Glide is a wall-directed swim toward the maze centre, typically initiating a trial from the start arm along the arm wall. Sharp Turn is a goal-oriented turnaround at or near the maze centre, where the mouse pivots toward the target arm (West) that holds the escape platform. Straight Sprint is a direct, momentum-preserving swim along the long axis of the target arm. Platform is platform climbing and was excluded from downstream sequence analyses, since its kinematics encode post-escape motor activity and no navigational content.

Pose-anchored skeleton strips illustrate the characteristic postural trajectory of each syllable (Fig. 4c). Bout centroid paths pooled across early training (Tr1 and Tr2; Fig. 5b) and late training plus the pre-anaesthesia test (Tr5 and Test; Fig. 5c) show how the spatial deployment of each syllable sharpens with task acquisition: Wall Turn and Intermediate Swim saturate the arm peripheries early in training, whereas by late training, Straight Sprint concentrates along the long axis of the target arm and Sharp Turn clusters near the maze centre. A representative learned-sequence schematic captures the Wall Glide → Sharp Turn → Straight Sprint → Platform chain that defines a successful escape in a well-trained animal (Fig. 5d).

**Figure 5:**
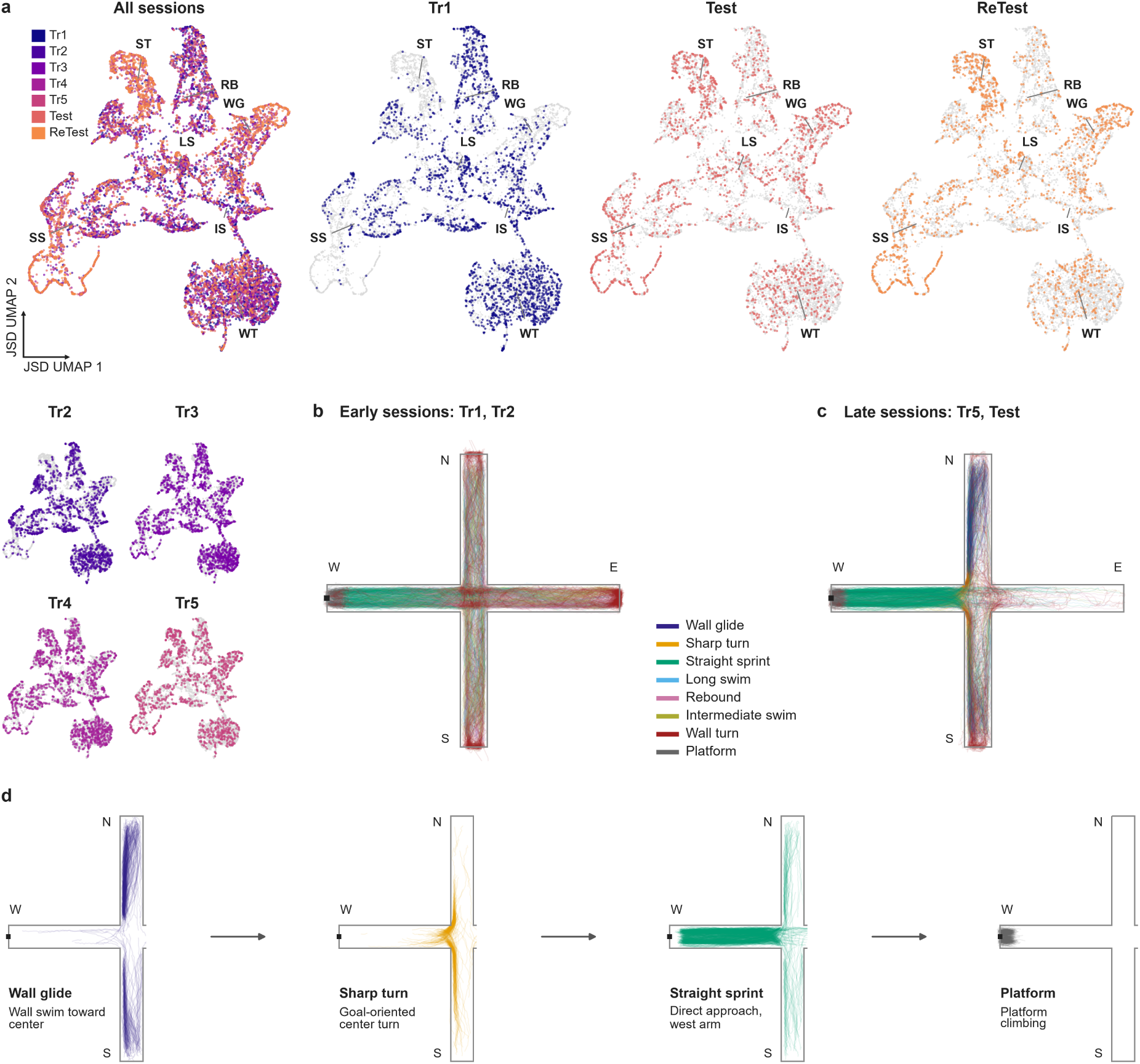
Syllable use and spatial deployment sharpen with learning in the Water Cross Maze. (a) Uniform manifold approximation and projection (UMAP) embedding of the 12,053 bouts (42 mice, LA n = 24, HA n = 18) from the Jensen-Shannon distances between their merged Gibbs marginal probability vectors (the embedding of Figure 4a), by session: all seven sessions overlaid (five training sessions Tr1-Tr5, pre-anaesthesia Test and post-anaesthesia ReTest), then Tr1, Test and ReTest, and Tr2-Tr5, each with that session’s bouts in colour over all bouts in grey. Syllable abbreviations as in Figure 4c; Platform has no bouts in this embedding. (b) Centroid paths of the 4,213 bouts of the early training sessions (Tr1, Tr2, 42 mice) on the Water Cross Maze, one line per bout, coloured by syllable; N, S, W, E, the maze arms; black square, the end of the west arm. (c) As (b) for the 3,661 bouts of the late sessions (Tr5, Test, 42 mice). (d) The four-step escape sequence Wall glide, Sharp turn, Straight sprint, Platform: each step shown as the late-session bout paths of that syllable (2,091 bouts, all 42 mice in each step) on the north, south and west arms, with a short description of the step; arrows, order of the steps.

Two independent projections corroborated the 8-syllable decomposition. A uniform manifold approximation and projection (UMAP) built on the Jensen-Shannon distance (JSD) across bout-level syllable-composition vectors produced seven well-separated clusters (Fig. 4a; UMAP geometry is shown for within-cluster dispersion only and between-cluster distances should not be read as quantitative). A second UMAP was built directly from 15 pose-derived kinematic features to test whether the syllables also differ kinematically, and 4 of the 8 syllables were recovered (Sharp Turn, Straight Sprint, Wall Turn and Platform). The four unrecovered ones (Wall Glide, Long Swim, Rebound and Intermediate Swim) were kinematically similar swimming variants that the model separates through temporal context (Fig. 4b,d; Supplementary Fig. 6b). The eight syllables therefore formed a reproducible vocabulary whose spatial deployment sharpened with learning, and every trial could be read as a sequence of these syllables.

### Post-anaesthetic navigation sequences

Syllable usage and syllable sequence were then compared between the phenotypes across training, Test and ReTest. Repertoire composition did not separate the groups at any timepoint. Shannon entropy of the 7-syllable frequency vector was comparable between LA and HA across training, Test, and ReTest (all p > 0.36; Supplementary Fig. 8f). Principal component analysis and UMAP of the frequency vectors placed LA and HA mice in overlapping distributions at every test day, and the JSD between median group profiles was similar at Test (0.140) and ReTest (0.144). Frequency did shift across learning, with Intermediate Swim and Rebound declining while Wall Glide and Straight Sprint increased. These changes tracked task acquisition and were unchanged by anaesthetic exposure (Fig. 5a, Fig. 6a,b).

**Figure 6:**
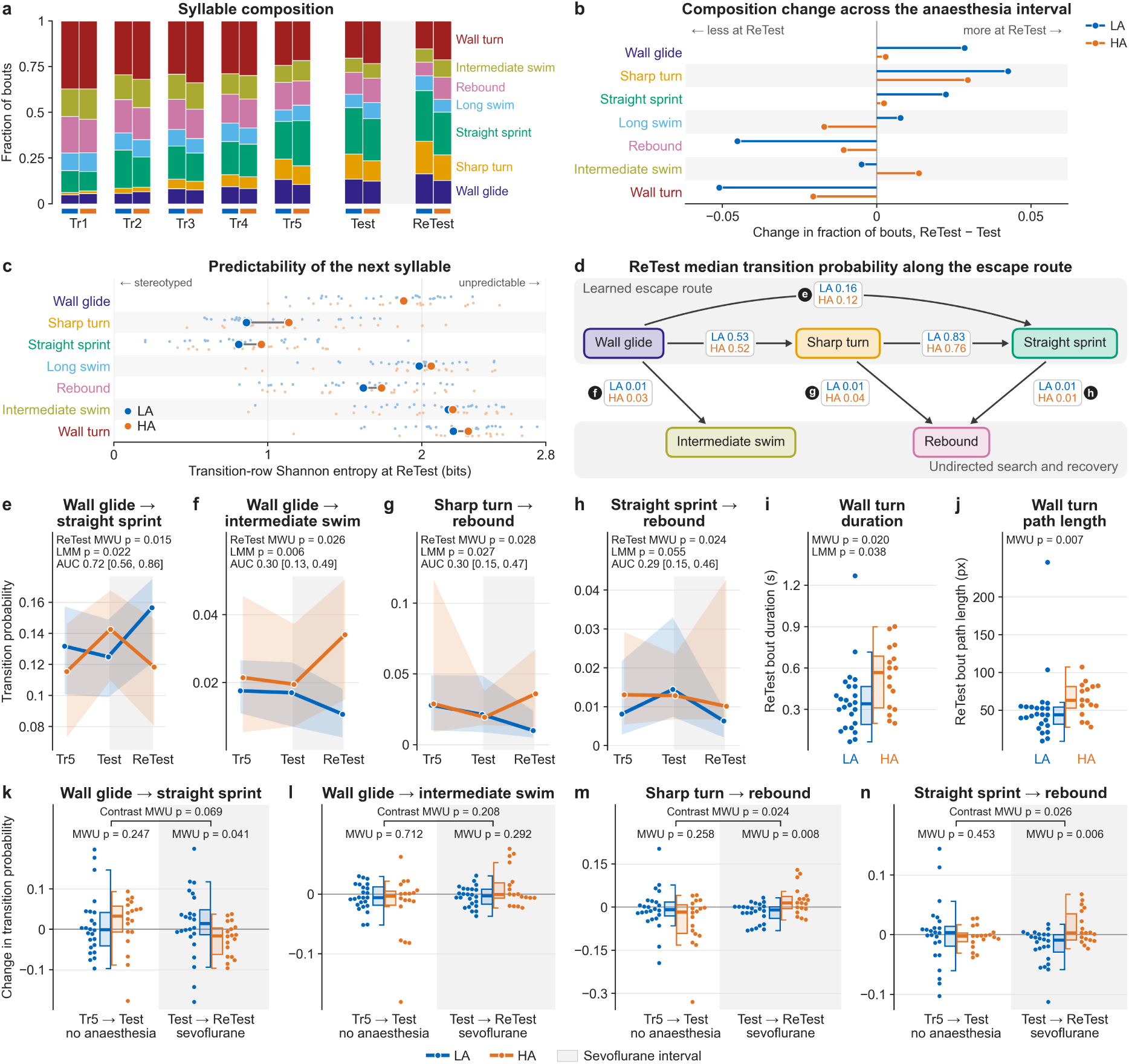
Sevoflurane disrupts the learned navigation sequence in HA mice while the syllable repertoire stays unchanged. (a) Syllable frequency per session (fractional counting), stacked per group, across the five training days (Tr1 to Tr5), the pre-anaesthesia Test and the post-anaesthesia ReTest for low anxiety (LA, blue, n = 24) and high anxiety (HA, orange, n = 18) mice; LA and HA bars side by side per session, colour strip under each bar, group; syllables named beside the ReTest bars. (b) Change in each syllable’s frequency from Test to ReTest (ReTest minus Test), LA above HA in each row (LA n = 24, HA n = 18); stems from zero. (c) Shannon entropy of the transition row of each syllable at ReTest, in bits, from 0 to log2(7) (LA n = 24, HA n = 18). (d) Group median conditional transition probability at ReTest between Wall Glide, Sharp Turn, Straight Sprint, Intermediate Swim and Rebound, printed per link (LA above HA, LA n = 24, HA n = 18); circled letters mark the transitions shown in (e-h) and (k-n). (e-h) Conditional transition probability at Tr5, Test and ReTest for Wall Glide → Straight Sprint (e), Wall Glide → Intermediate Swim (f), Sharp Turn → Rebound (g) and Straight Sprint → Rebound (h); bold lines, group medians; bands, 95% bootstrap confidence interval of the median (10,000 resamples of mice); MWU p, LMM p and AUC with 95% bootstrap CI at ReTest, above 0.5 larger in LA. (i) Wall Turn bout duration in s and (j) Wall Turn bout path length in px at ReTest (LA n = 24, HA n = 16). (k-n) Within-mouse change in the transition probabilities of (e-h) over the interval without anaesthesia (Tr5 → Test) and the interval with sevoflurane (Test → ReTest) in LA (n = 24) and HA (n = 18) mice, MWU p per interval; contrast, MWU p on each mouse’s change with sevoflurane minus its change without. Grey, interval with sevoflurane (in a, the gap between the Test and ReTest bars). MWU, two-sided Mann-Whitney U test, uncorrected; LMM, linear mixed model with batch as random intercept. Points, individual mice (faint in c); large filled circles in c, group medians; half-boxes, median and interquartile range, whiskers to the most extreme observation within 1.5 interquartile ranges. P values are printed in the panels, and no significance symbols are used.

But anaesthesia changed how those syllables were linked to each other: the transition matrix separated LA from HA across the post-anaesthetic tests. Across Tr1 to Tr5, the entropy rate of the bout-level syllable chain fell from 1.88 to 1.57 bits and remained low at ReTest (1.51 bits), indicating preserved sequence structure in both groups (Supplementary Fig. 8g). The full transition matrices at Test and ReTest and their ReTest group difference showed a preserved syllable sequence in LA mice and a diffuse redistribution in HA mice, concentrated in the cells connecting Sharp Turn, Straight Sprint, Wall Glide, and Rebound (Supplementary Fig. 8a to e). At ReTest the whole-matrix transition distance did not separate the groups (MWU p = 0.199), confirming that the divergence lay in individual transitions while the matrix stayed stable. Four ReTest cells converged on a single direction (Fig. 6d to h). The learned Wall Glide → Straight Sprint transition (priming to approach) was maintained in LA and lost in HA (MWU p = 0.015; Fig. 6e). Three further cells captured a diversion into undirected searching in HA: Sharp Turn → Rebound (MWU p = 0.028; Fig. 6g), Straight Sprint → Rebound (MWU p = 0.024; Fig. 6h), and Wall Glide → Intermediate Swim (MWU p = 0.026; Fig. 6f). Three of the four cells held after adjustment for batch as a random effect. The fourth (Straight Sprint → Rebound) was marginal, with an effect size indistinguishable from the other three. No single cell survived false discovery rate correction at the per-cell level of the 7 × 7 matrix (0 of 49 at ReTest, 0 of 49 at Test).

The four transitions that differed between LA and HA mice at ReTest were also compared over two 3-day intervals: the interval without anaesthesia (Tr5 to Test) and the interval with sevoflurane (Test to ReTest). None of the four transitions differed significantly between LA and HA mice at Tr5 or at Test (Fig. 6e to h). Over the interval without anaesthesia, the change in these transitions did not differ significantly between LA and HA mice (Fig. 6k to n). Over the interval with sevoflurane, the change in three of the four transitions differed between LA and HA mice: Wall Glide → Straight Sprint (MWU p = 0.041; Fig. 6k), Sharp Turn → Rebound (p = 0.008; Fig. 6m), and Straight Sprint → Rebound (p = 0.006; Fig. 6n). The median change from Test to ReTest was positive in LA mice and negative in HA mice for Wall Glide → Straight Sprint. For Sharp Turn → Rebound and Straight Sprint → Rebound, the median change was positive in HA mice and negative in LA mice. The change in Wall Glide → Intermediate Swim over this interval did not differ significantly between LA and HA mice (Fig. 6l). The difference between the two interval changes of each mouse (the change with sevoflurane minus the change without anaesthesia) was nominally larger in HA than in LA mice for Sharp Turn → Rebound (p = 0.024; Fig. 6m) and Straight Sprint → Rebound (p = 0.026; Fig. 6n). The same difference did not reach significance for Wall Glide → Straight Sprint or Wall Glide → Intermediate Swim (Fig. 6k,l). In these four transitions, a difference between LA and HA mice was detected only at ReTest, the session after the interval with sevoflurane.

During training, sequence learning was driven almost entirely by Sharp Turn, whose row entropy dropped 0.79 bits between Tr1 and Tr5. At ReTest, the LA−HA difference localized to exactly the three source syllables implicated by the transition-cell analysis: Sharp Turn (+0.28 bits HA minus LA), Straight Sprint (+0.15 bits), and Rebound (+0.12 bits) (Fig. 6c). The pattern was reversed or absent at Test (Sharp Turn −0.19 bits; Straight Sprint −0.18 bits). After sevoflurane, transitions out of these key syllables became less predictable in HA mice while remaining ordered in LA mice, and this localization within the chain did not depend on any single per-cell test.

Swim speed during these bouts was comparable between groups (MWU p = 0.384), whereas HA mice spent significantly longer in the Wall Turn syllable, with longer bout duration (MWU p = 0.020) and greater path length (MWU p = 0.007; Fig. 6i,j). These bout kinematics are from the 24 LA and 16 HA mice with Wall Turn bouts at ReTest, since two HA mice had none. The duration difference held after adjustment for batch, and the path-length difference did not. Wall Turn frequency itself trended in the same direction without reaching significance (MWU p = 0.151), consistent with the unchanged repertoire composition. Together, the transition-cell and per-row entropy analyses localized the effect within the syllable chain: sevoflurane selectively disrupted the learned sequential organization of spatial navigation in HA mice, an effect invisible to conventional scoring and to repertoire measures.

### Baseline sleep/wake architecture

Sleep at baseline was compared first, to establish whether the two phenotypes already differed before any anaesthetic. Baseline EEG/electromyography was recorded for 24 h on D1, two weeks before sevoflurane exposure, and yielded complete 24 h recordings in 38 of 42 mice (22 LA, 16 HA; Supplementary Fig. 9a). Vigilance states were analysed separately by light phase (Zeitgeber time, ZT; ZT12 to ZT24) and dark phase (ZT0 to ZT12, active). Dark-phase vigilance-state proportions did not significantly differ between the two groups. The largest difference was in dark-phase REMS, higher in HA mice, though it did not reach significance (median 2.04% vs 1.28%; MWU p = 0.204; Supplementary Fig. 9b,c). Dark-phase NREMS was likewise higher and dark-phase WAKE lower in HA mice, without reaching significance (NREMS MWU p = 0.084; WAKE MWU p = 0.074). Light-phase proportions were indistinguishable across all three vigilance states. LA mice had shorter NREMS latency than HA mice at the start of the light phase, though the difference was marginal (NREMS latency median 8 s vs 16 s; MWU p = 0.050; Supplementary Fig. 9j). Spectral features separated LA and HA during WAKE but not during sleep. Dark-phase WAKE alpha power (8 to 13 Hz) was higher in LA mice (MWU p = 0.012; Supplementary Fig. 9h,i). Light-phase WAKE theta power (4 to 8 Hz) showed the same direction but did not reach significance (MWU p = 0.060). NREMS and REMS spectra did not differ in any of the five frequency bands. Aperiodic parameters were equivalent across groups, with a single isolated light-NREMS offset difference (MWU p = 0.043; Supplementary Fig. 9n).

Sleep spindles added one dark-phase asymmetry. HA mice produced more spindles per 12 h dark phase than LA mice at baseline (median 37.75 vs 29.75; MWU p = 0.036; Supplementary Fig. 9d), while spindle density, duration, and amplitude were equivalent (Supplementary Fig. 9e to g). Microarchitecture outside these targeted measures showed no group differences. Bout length did not differ between LA and HA in any state or phase (all MWU p > 0.36; Source Data), and Markov vigilance-state transitions and PeEn were indistinguishable between LA and HA across every state and phase tested (Supplementary Fig. 9l,m). At baseline, the phenotype therefore showed in the active phase as higher WAKE alpha power in LA mice and more spindles in HA mice, while sleep/wake amounts and sleep microarchitecture were comparable between groups.

### Post-anaesthetic REMS response

Finally, the sleep/wake response to sevoflurane was examined in the two phenotypes. Post-anaesthetic sleep on D18 (Fig. 7a to f) was compared to each animal’s own pre-anaesthetic baseline (D1) as a within-subject change (post general anaesthesia minus baseline). Thirty-one mice contributed complete paired recordings (19 LA, 12 HA). Because baseline sleep values differed modestly between groups, every change metric was also fit with a baseline-adjusted LMM.

**Figure 7:**
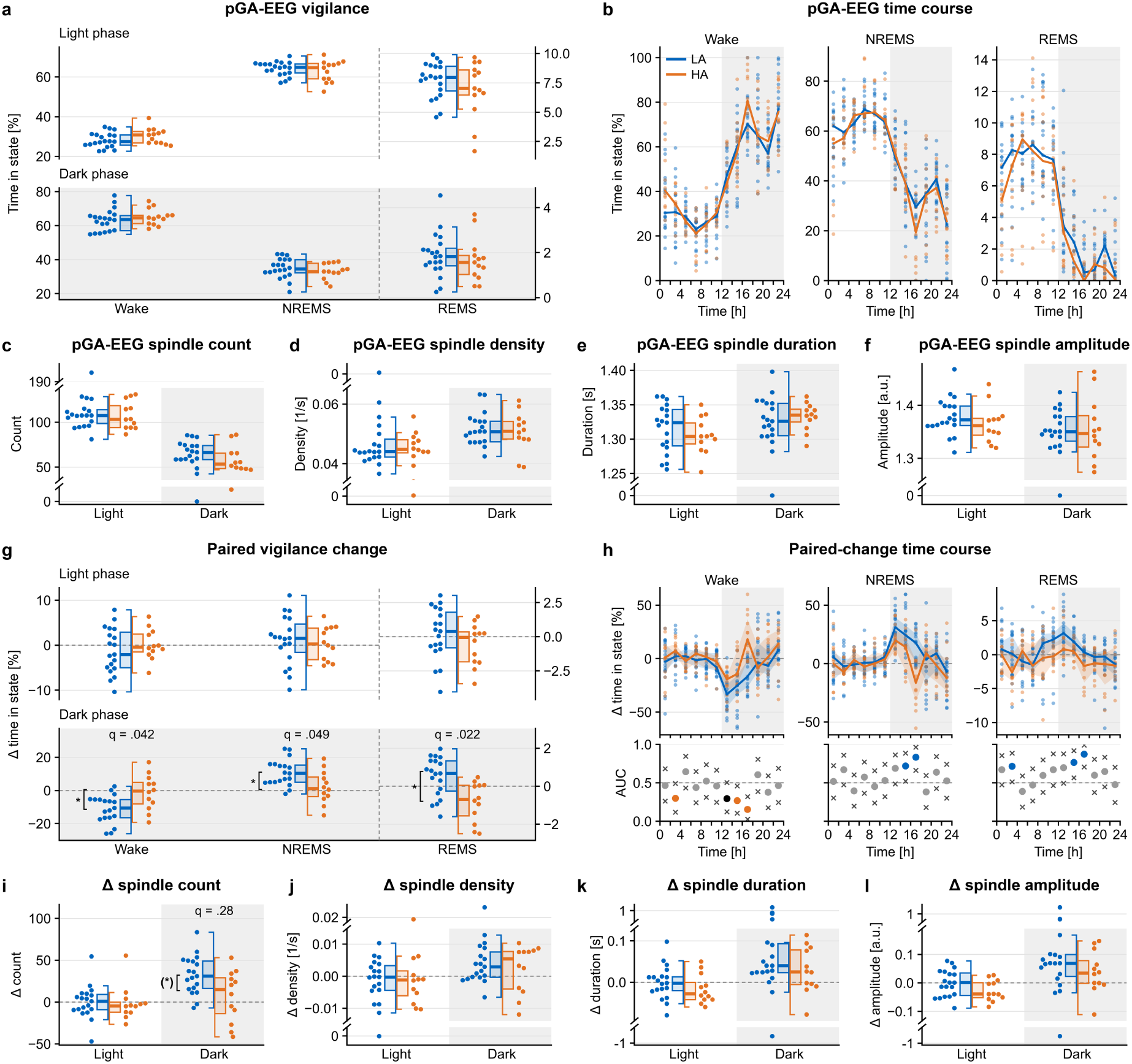
LA and HA mice show opposing dark-phase rapid eye movement sleep (REMS) responses to sevoflurane. (a) Percentage of time spent in WAKE, non-rapid eye movement sleep (NREMS) and REMS during the light phase and the dark phase of the post-anaesthetic 24-h recording (pGA-EEG; LA n = 20, HA n = 13); REMS on its own right-hand scale. (b) Vigilance state percentages in 2-hour bins across the post-anaesthetic recording, one panel per state: faint points, individual mice (n = 31, LA n = 19, HA n = 12, mice with both baseline and post-anaesthetic recordings); lines, group medians; dark phase shaded. (c-f) Post-anaesthetic sleep spindle metrics per phase (LA n = 19, HA n = 12): (c) spindle count, (d) spindle density, (e) median spindle duration, (f) median normalized amplitude. (g) Within-subject change scores (Δ = pGA − bEEG, the post-anaesthetic recording minus the baseline recording) in the percentage of time in each state per phase (n = 31; LA n = 19, HA n = 12); dashed line, Δ = 0. (h) Within-subject change in vigilance state percentages per 2-hour bin: line, group median; band, 95% CI (bootstrap of the median, 10,000 resamples of mice); faint points, individual mice (LA n = 19, HA n = 12, HA n = 11 in the bin centred at 23 h). Lower strip, AUC (LA > HA) per bin as circles with 95% CI (bootstrap, 2,000 resamples) as crosses; dashed line, AUC = 0.5. AUC dots: grey, 0.3-0.7; outside this range, blue or orange when the CI excludes 0.5, otherwise black. (i-l) Within-subject change in (i) spindle count, (j) spindle density, (k) median spindle duration, (l) median normalized amplitude (LA n = 19, HA n = 12). (a, c-g, i-l) Dots, individual mice; half-boxes, median and interquartile range, whiskers to the most extreme observation within 1.5 interquartile ranges; observations beyond the axis break are drawn in the stub above or below it. q, Benjamini-Hochberg FDR-corrected p of the two-sided Mann-Whitney U test; bracket with asterisk, q < 0.05; bracket with (*), uncorrected p < 0.05 only.

The primary finding was a reversal of direction in dark-phase REMS. LA mice showed a REMS rebound after sevoflurane while HA mice lost REMS (LA median change +0.68 percentage points, HA −0.69; LMM p = 0.007; Fig. 7g,h). The REMS reversal was accompanied by a larger dark-phase NREMS gain in LA than in HA mice (LA +10.3 percentage points, HA +1.2; LMM p = 0.007) and a dark-phase WAKE decrease confined to LA mice (LA −10.6 percentage points, HA −0.35; LMM p = 0.004). These three changes form a single redistribution during the dark phase. LA mice converted ∼10 percentage points of dark-phase WAKE into sleep, split between NREMS and a smaller REMS rebound, whereas HA mice remained near their baseline. Light-phase changes were weaker: no light-phase change reached significance (REMS MWU p = 0.25, WAKE MWU p = 0.45), and neither light-phase REMS nor WAKE survived baseline adjustment. Dark-phase spindle count increased more in LA than HA (LA median +31, HA +15; LMM p = 0.022; Fig. 7i), with no change in density, duration, or amplitude (Fig. 7j to l). After adjustment for baseline, the dark-phase NREMS and WAKE changes weakened and no longer reached significance, while the unadjusted group differences remained significant (NREMS MWU p = 0.024, WAKE MWU p = 0.014). No standard error or p value could be estimated for the baseline-adjusted model of the dark-phase REMS change (its random-effects term fell to zero), so the REMS change is reported from the unadjusted comparison (LMM p = 0.007). The remaining change comparisons were not significant. Baseline-adjusted LMM estimates for every stage and phase are shown in Supplementary Fig. 9k. The direction of the sleep/wake response to sevoflurane therefore depended on the pre-anaesthetic phenotype. With the intra-anaesthetic EEG and the learned navigation sequence, sleep was the third outcome domain in which one label defined before induction tracked the response to the same exposure.

## Discussion

The brain’s response to sevoflurane during anaesthesia and recovery differed according to an anxiety phenotype defined before induction. At Adequate anaesthetic depth, high-anxiety (HA) mice showed relatively more high-frequency power in the aperiodic EEG background than low-anxiety (LA) mice under comparable measured exposure.

After anaesthesia, HA mice differed from LA mice in how particular behavioural patterns were linked within the learned navigation sequence, even though choice accuracy remained high and the escape-latency difference was small. The same phenotype also distinguished the sleep/wake response: dark-phase rapid eye movement sleep (REMS) increased in LA mice and decreased in HA mice relative to the pre-anaesthetic baseline. This combination of pre-anaesthetic phenotyping and exposure without concurrent surgery establishes a mouse model for directly testing the contribution of anxiety-related circuitry to anaesthetic vulnerability and interventions intended to improve recovery.

The greater relative high-frequency power in the aperiodic EEG background of HA mice suggests an anxiety-related difference in cortical state within broadly comparable measured anaesthetic conditions. In a model of population electrical activity, a flatter background spectrum arises when fast excitatory currents contribute more strongly relative to slower inhibitory currents ^61^. During propofol-induced loss of responsiveness in humans, the background spectrum became steeper ^62^. Higher trait anxiety was also associated with a smaller negative EEG response to prefrontal magnetic stimulation in humans, interpreted as a marker of reduced cortical inhibition ^63^. Our group difference concerned the distribution of aperiodic power, while the fitted exponent describing spectral steepness did not differ significantly. We hypothesize that sevoflurane reveals an anxiety-related difference in cortical excitability, with weaker inhibition in HA mice at the depth scored as Adequate in both groups ^61, 63^.

Preserved choice accuracy after sevoflurane anaesthesia coincided with longer Wall Turn bouts in HA mice and differences between LA and HA in selected syllable transitions. Longer Wall Turn bouts could delay arrival at the correct platform and contribute to the small escape-latency difference in HA mice despite comparable speed within these bouts. Differences in the organization of the learned navigation sequence were also observed at specific syllable transitions in exploratory analyses. While only time passed between the last training day and Test, LA and HA mice did not differ in how these transitions changed. When the sevoflurane exposure was added to an interval of the same length, LA and HA mice diverged in these transitions by ReTest. We therefore hypothesize that sevoflurane reorganized the learned navigation sequence in an anxiety-dependent manner. Studies of pain and associative learning in mice have shown how detailed behavioural profiling reveals differences that conventional summary measures do not capture ^64, 65^. Analyses of behavioural transitions have further distinguished phenotypes through the temporal organization of behaviour ^66, 67^. We extended earlier analyses of transitions between assigned syllables ^68, 69^ by retaining probabilities across possible assignments using the Gibbs marginals available in Keypoint-MoSeq ^41^. To our knowledge, the present study is the first to use probabilistic syllable assignments from Gibbs marginals in behavioural analyses, incorporating uncertainty in syllable assignment into comparisons between anxiety phenotypes.

After sevoflurane, dark-phase REMS rose in LA mice and fell in HA mice compared to their own baselines. The LA increase is consistent with the REMS rebound reported after volatile anaesthetics ^18, 34, 35^. This increase compensates for REMS lost during anaesthesia and would be expected in both groups, whereas HA mice showed a decrease. Before sevoflurane sleep/wake amounts were comparable between LA and HA mice, but HA mice had more dark-phase sleep spindles and LA mice higher dark-phase WAKE alpha power. A correlate of the anxiety phenotype was therefore present in these two dark-phase sleep/wake measures before sevoflurane. In rodents, uncontrollable stress and fear reminders reduce REMS through amygdala-dependent circuits, and the reduction is larger in the more stress-reactive BALB/cJ strain than in C57BL/6J mice ^70, 71^. In mice, thirty minutes of sevoflurane raised serum corticosterone and produced a post-anaesthesia stress response that depended on hypothalamic corticotropin-releasing hormone neurons ^72^. We hypothesize that a larger stress response to sevoflurane and to the ReTest on the following day produced a REMS reduction in HA mice that exceeded the compensatory increase. If so, the direction of the post-anaesthetic REMS change depends on the size of the stress response, since the anaesthetic exposure was matched between groups.

If the cortical difference under sevoflurane reflects an anxiety-related difference in cortical excitability, the same difference could underlie the navigation and REMS responses. Two systems that set cortical state under anaesthesia also act on REMS. In rats, locus coeruleus noradrenergic neurons discharge most during waking and fall almost silent during REMS ^73^, and their activation reduced delta power and accelerated emergence during continuous isoflurane anaesthesia ^74^. Corticosterone altered cortical inhibition through GABAergic and purinergic signalling in acute mouse brain slices ^75^. We hypothesize that arousal and stress systems are more active in HA mice. Under this hypothesis, one difference produces more high-frequency aperiodic EEG power at Adequate depth, differences in selected transitions of the learned navigation sequence after anaesthesia, and a REMS decrease larger than the compensatory increase. If so, the anxiety phenotype marks the activity of arousal and stress systems, which sets the response to the anaesthetic before induction.

Both measures linked in the mouse model are available in patients, namely the anxiety before induction and the aperiodic EEG during maintenance. In a systematic review of older surgical patients, POA was associated with postoperative delirium when anxiety was dichotomized using a threshold on an anxiety scale ^9^. No association was found when anxiety was analysed as a continuous score ^9^. This threshold pattern is consistent with a phenotype, as in the LA and HA grouping of the mice. In surgical patients, the aperiodic component and its SEF95 can be extracted from frontal EEG recordings ^56^. Delirium in the recovery unit was more frequent in patients without the spindle-dominant anaesthesia pattern in the emergence EEG, and readmission or a hospital stay of more than six days was more common after such delirium ^76^. In mice, learning and memory were unimpaired after isoflurane alone and impaired after surgery under isoflurane ^15^. In the present study, conventional maze scoring after sevoflurane alone was likewise a negative finding, with no difference between the phenotypes. By contrast, the organization of learned behaviour differed between LA and HA mice after the same exposure. Reaching the platform is a simple endpoint that HA mice still met after sevoflurane. The effect of the anaesthetic was seen in the way the platform was reached. In patients, the open question raised by these findings is whether POA is expressed in the EEG at Adequate anaesthetic depth, and whether that intra-anaesthetic signature is associated with delirium and later recovery beyond the anxiety score itself.

## Conclusion

LA and HA mice responded differently to the same sevoflurane exposure: in the EEG at Adequate anaesthetic depth, and afterwards in the organization of learned navigation and in sleep. All three responses were separated by the anxiety phenotype that had been defined before any anaesthetic. Vulnerability to an anaesthetic therefore differs between individuals and is identifiable before induction. Because sevoflurane was given without surgery, each of these responses is attributable to the anaesthetic alone. The next step in this model is a chemogenetic manipulation of the proposed arousal and stress systems before the anaesthetic is given, with the effects measured on the EEG during anaesthesia, on learned behaviour and on sleep in the same animals. In learned behaviour, the effect of sevoflurane appeared in the organization of the navigation sequence while the platform was still reached. After anaesthesia, how a learned task is performed is therefore an outcome measure alongside whether the task is solved. Anxiety before induction is measurable in every surgical patient, and the mechanisms by which the brain’s response to an anaesthetic differs with this trait can now be tested in mice.

## Data availability

The processed data needed to reproduce the results will be available at Zenodo (https://doi.org/10.5281/zenodo.23995877) upon publication. The deposit comprises the DeepLabCut keypoints, the fitted Keypoint-MoSeq model with its Gibbs marginal probabilities, the scored EEG and EMG recordings with their hypnograms, the intra-anaesthetic EEG recordings, the fear conditioning and Water Cross Maze scores, and the group assignments. Source data are provided with this paper. Raw video recordings and raw EEG recordings are available from the corresponding author on request.

## Code availability

The custom Python and MATLAB code for video preprocessing, pose estimation, Keypoint-MoSeq model fitting, behavioural syllable analysis, EEG analysis, statistics and figure generation will be available at Zenodo (https://doi.org/10.5281/zenodo.23995680) under the MIT licence upon publication.

## Acknowledgements

The authors thank Andreas Blaschke for technical support and Stefanie Monecke for animal husbandry and animal management.

## Author contributions

AA: Experimental design, Investigation, Formal analysis, Writing, Editing. MV: Investigation. AK: Investigation, Formal analysis. CH: Investigation, Formal analysis. SW: Formal analysis, Data curation. MVS: Methodology, Review. MK: Methodology. GR: Methodology, Review. GS: Review, Financial support. TF: Experimental design, Conceptualization, Methodology, Writing, Review, Supervision. All authors approved the final version of the manuscript.

