## Supporting Information for "Pre-anaesthetic anxiety phenotype stratifies anaesthesia EEG, acquired navigation, and post-anaesthesia sleep in sevoflurane-exposed mice"

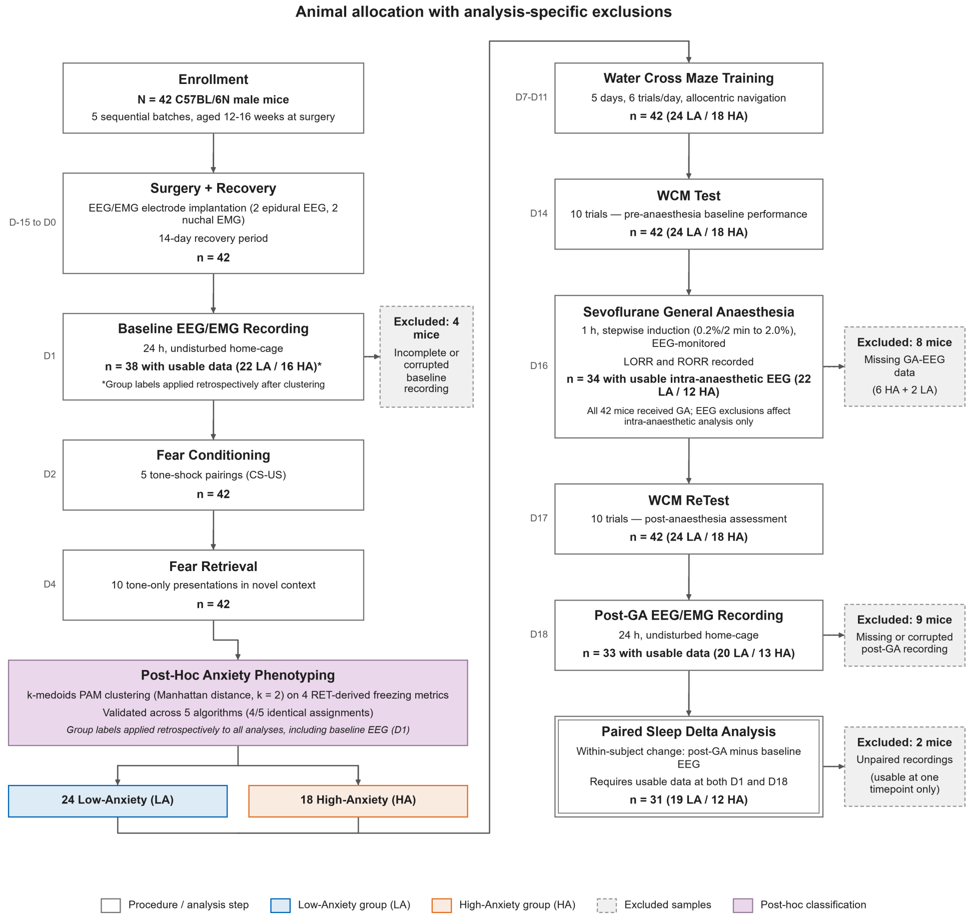


**Supplementary Figure 1:** Animal allocation with analysis-specific exclusions. Flow diagram of the experimental procedures, anxiety phenotyping and samples contributing to the EEG

and behavioural analyses, with the number of mice (n) and the exclusions at each step given in the boxes. LA, low anxiety; HA, high anxiety.

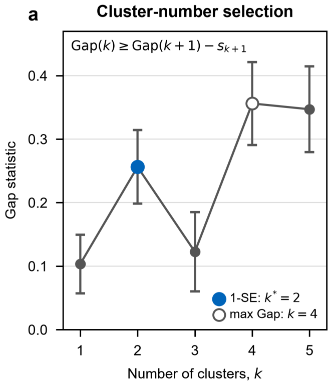


**b** **Algorithm sensitivity (fixed k = 2)**

| Method | Split LA/HA | Match to primary | Mean silhouette |
|---|---|---|---|
| **Manhattan K-med.** | 24/18 | — | 0.428 |
| Euclidean K-med. | 24/18 | 42/42 | 0.417 |
| K-means | 24/18 | 42/42 | 0.417 |
| Ward | 24/18 | 42/42 | 0.417 |
| Full-cov. GMM | 22/20 | 40/42 | 0.387 |

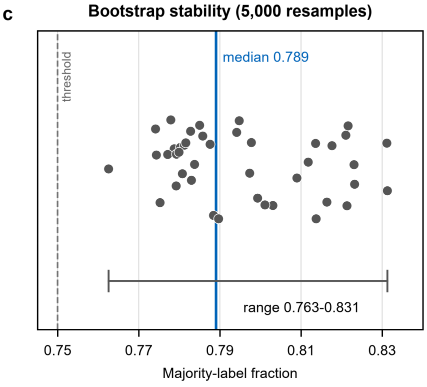


**Supplementary Figure 2:** Internal validation of the k-medoids clustering. (a) Gap statistic for k = 1 to 5 clusters (k-medoids clustering, Manhattan distance, 5,000 Monte Carlo references); error bars, standard error $s_k$. Filled blue circle, k* = 2 selected by the 1-SE rule Gap(k) ≥ Gap(k + 1) − $s_{k+1}$; open circle, maximum gap at k = 4. (b) Algorithm sensitivity at fixed k = 2: LA/HA split, agreement with the primary k-medoids clustering (Manhattan distance) and mean silhouette value for five algorithms (n = 42 mice). (c) Bootstrap stability (5,000 resamples): majority-label fraction of each of the 42 mice; blue line, median (0.789); horizontal bar, range (0.763-0.831); dashed line, 0.75.

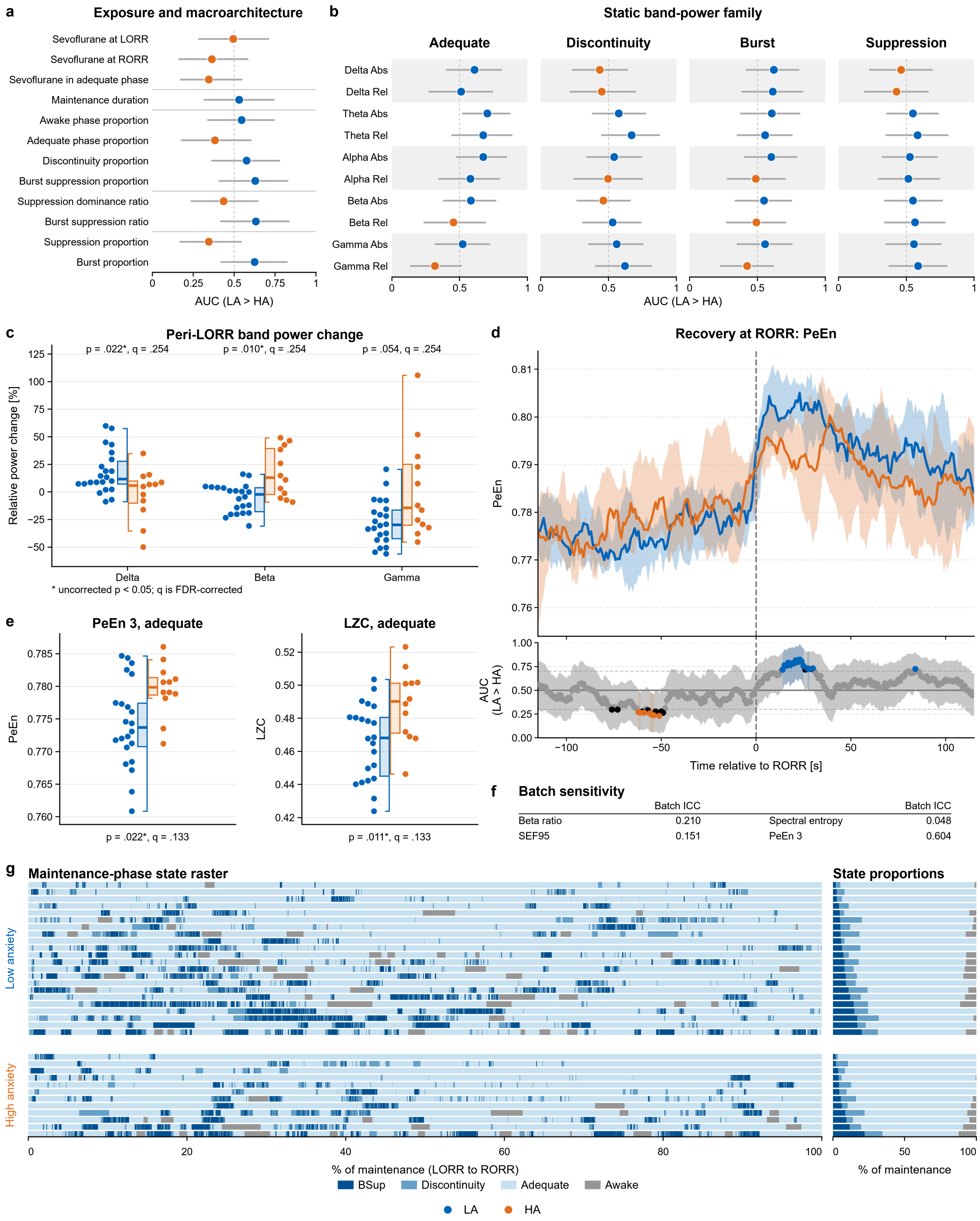


**Supplementary Figure 3:** Anaesthetic exposure, EEG macroarchitecture and spectral measures in LA and HA mice. (a) AUC (LA > HA) with 95% bootstrap CI for twelve anaesthetic-course measures, grouped as sevoflurane concentrations, maintenance duration, state proportions of maintenance, burst suppression ratios and burst suppression architecture; dashed line, AUC = 0.5. (b) AUC (LA > HA) with 95% bootstrap CI for absolute

(Abs) and relative (Rel) band power in five bands during four states; In (a, b), blue, AUC above 0.5; orange, below 0.5. (c) Relative band power change from before to after loss of righting reflex (LORR) in the delta, beta and gamma bands. (d) Permutation entropy (PeEn) around return of righting reflex (RORR; LA $n = 19$, HA $n = 8$): pointwise median and 95% CI (mouse-resample bootstrap, 10,000 resamples). Lower strip, pointwise AUC (LA > HA) with 95% CI; blue and orange dots, AUC outside 0.3-0.7 with CI excluding 0.5 (LA > HA and HA > LA, respectively); black dots, AUC outside 0.3-0.7 with CI including 0.5; dashed lines, 0.3 and 0.7. (e) PeEn and Lempel-Ziv complexity (LZC) during Adequate anaesthesia. (c, e) Dots, individual mice (LA $n = 22$, HA $n = 12$); half-boxes, median and interquartile range, whiskers to the most extreme observation within 1.5 interquartile ranges; p, two-sided Mann-Whitney U test, asterisk $p < 0.05$; q, Benjamini-Hochberg FDR-corrected p. (f) Batch intraclass correlation coefficient (ICC) of four spectral measures from linear mixed models (metric ~ group + (1 | batch), five batches). (g) Scored state of each mouse across the maintenance phase from LORR to RORR (LA $n = 22$, HA $n = 12$), one row per mouse: BSup, burst suppression; right, proportion of maintenance in each state.

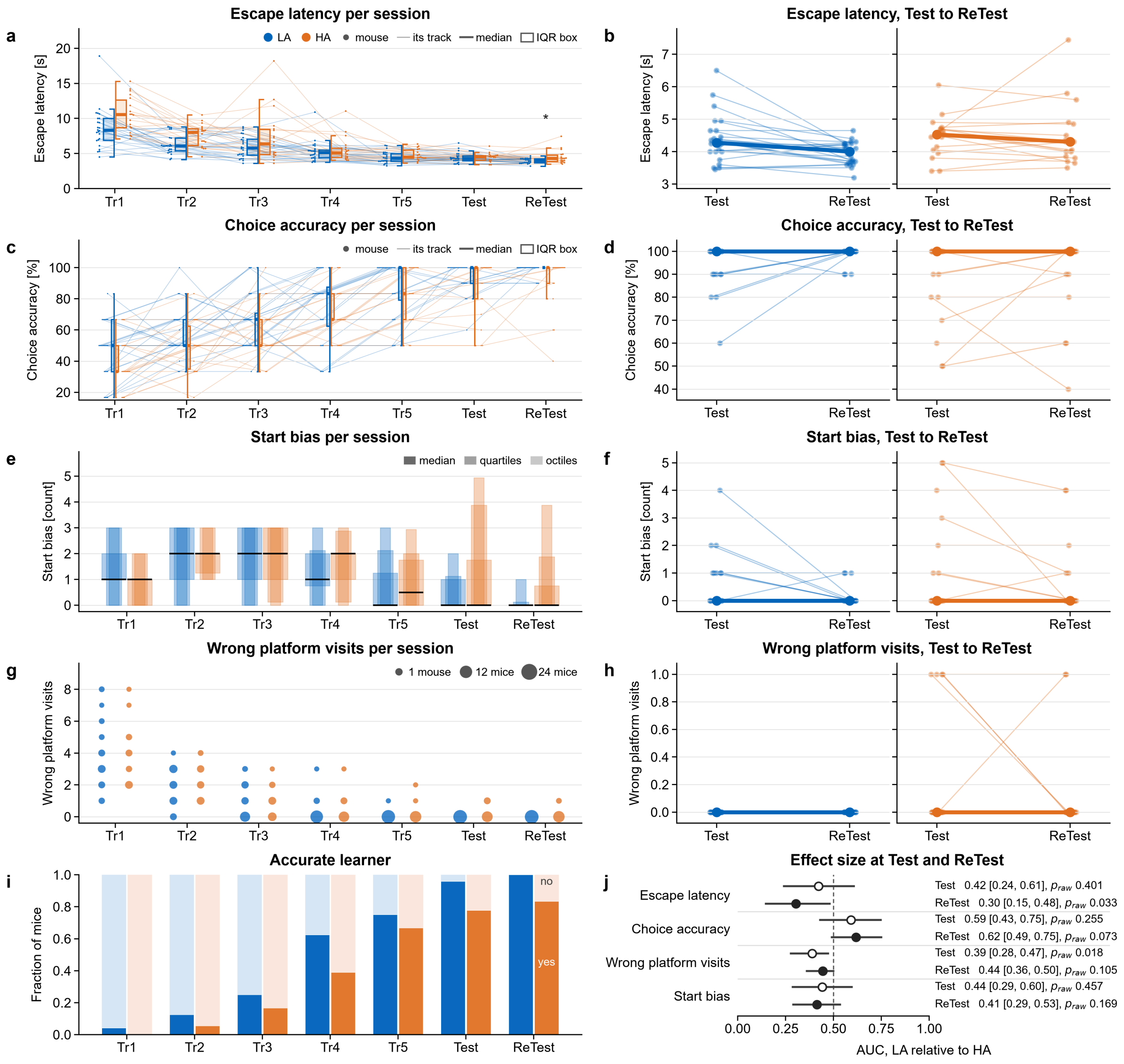


**Supplementary Figure 4:** Conventional scoring of Water Cross Maze performance across training, Test and ReTest in LA and HA mice. Sessions are the five training days (Tr1-Tr5), the pre-anaesthesia Test and the post-anaesthesia ReTest; low anxiety (LA, blue, n = 24) and high anxiety (HA, orange, n = 18). (a) Escape latency per session (asterisk, uncorrected two-sided Mann-Whitney U test p = 0.033 at ReTest). (b) Escape latency of each mouse from Test to ReTest. (c) Choice accuracy per session. (d) Choice accuracy of each mouse from Test to ReTest. (e) Start bias per session as letter-value boxes: black bar, median; darkest box, quartiles; middle box, octiles; palest box, sixteenths. (f) Start bias of each mouse from Test to ReTest. (g) Wrong platform visits per session; dot area, number of mice at that count (key: 1, 12 and 24 mice). (h) Wrong platform visits of each mouse from Test to

ReTest. (i) Fraction of mice in each group meeting the accurate learner criterion per session; yes, solid; no, pale. (j) AUC (LA relative to HA) with 95% bootstrap confidence interval for escape latency, choice accuracy, wrong platform visits and start bias at Test (open circles) and ReTest (filled circles); dashed line, AUC = 0.5; p raw, uncorrected p of the two-sided Mann-Whitney U test, printed beside each row. (a, c) Small points, individual mice; thin lines, each mouse across sessions; thick bar, group median; box, interquartile range. Upper whiskers are omitted where the 75th percentile equals 100% in (c). (b, d, f, h) LA left, HA right; thin lines, individual mice; thick line with circles, group medians.

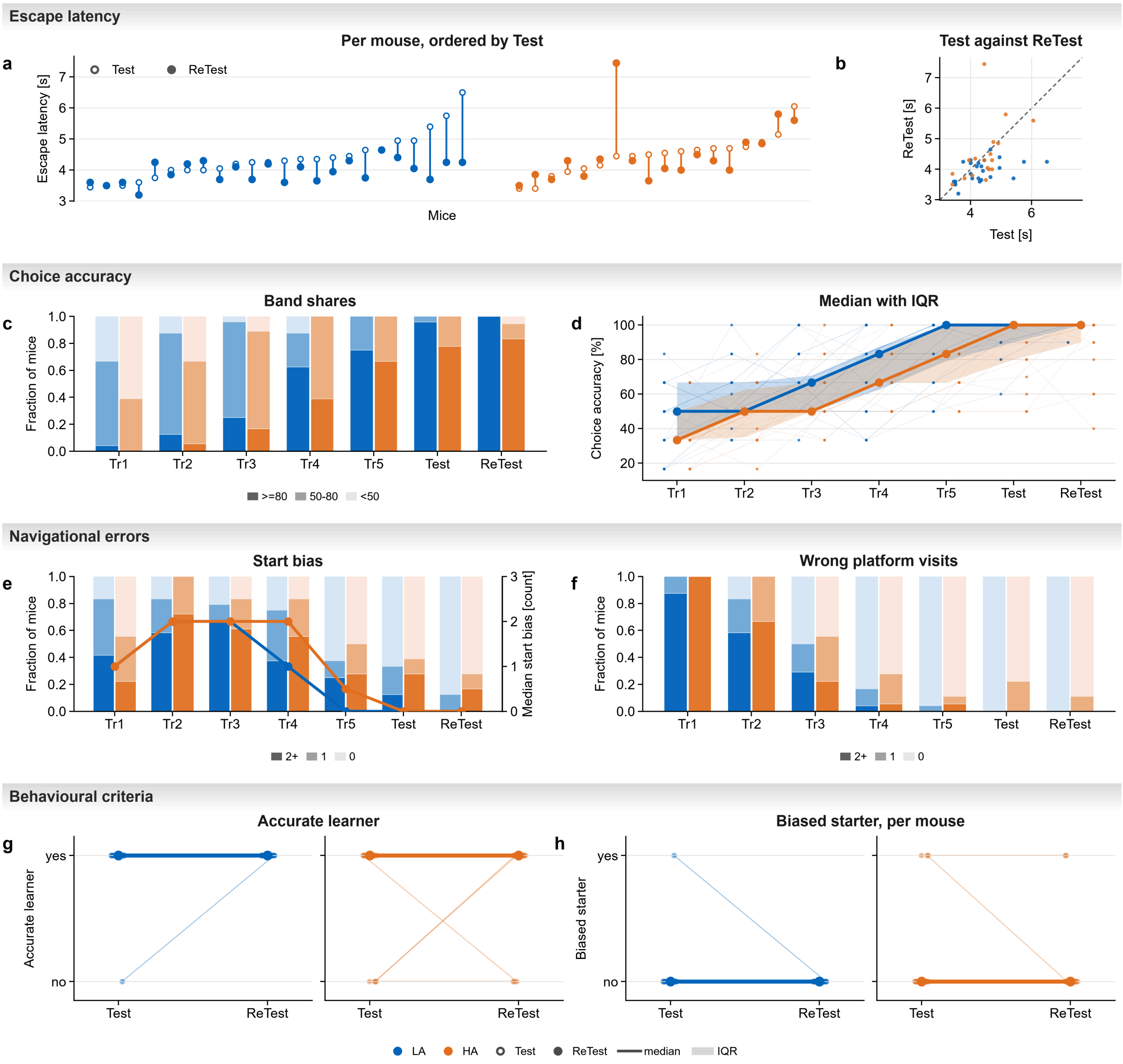


**Supplementary Figure 5:** Conventional Water Cross Maze metrics in LA and HA mice, per mouse and as group shares. Sessions are the five training days (Tr1-Tr5), the pre-anaesthesia Test and the post-anaesthesia ReTest; low anxiety (LA, blue, n = 24) and high anxiety (HA, orange, n = 18). (a) Escape latency of each mouse at Test and ReTest; mice ordered by their Test value within each group. (b) Escape latency at ReTest against Test. (c) Share of mice per session in three choice accuracy bands (≥80 %, 50-80 %, <50 %). (d) Choice accuracy per session. (e) Share of mice per session in three start bias bands (2 or more, 1, 0), with LA and HA median start bias counts across all seven sessions on the right-hand axis. (f) Share of mice per session in three wrong platform visit bands (2 or more, 1, 0). (g) Accurate learner status, yes or no, of each mouse from Test to ReTest. (h) Biased starter status, yes or no, of each mouse from Test to ReTest. Open and filled circles in (a), Test and ReTest, joined within mouse. Points in (b), individual mice; dashed line, identity. Faint points and lines in (d), individual mice; shaded bands, interquartile range across mice. Lines with filled circles in (d, e, g, h), group medians; thin lines in (g, h), individual mice. Stacked shades in (c, e, f), category shares.

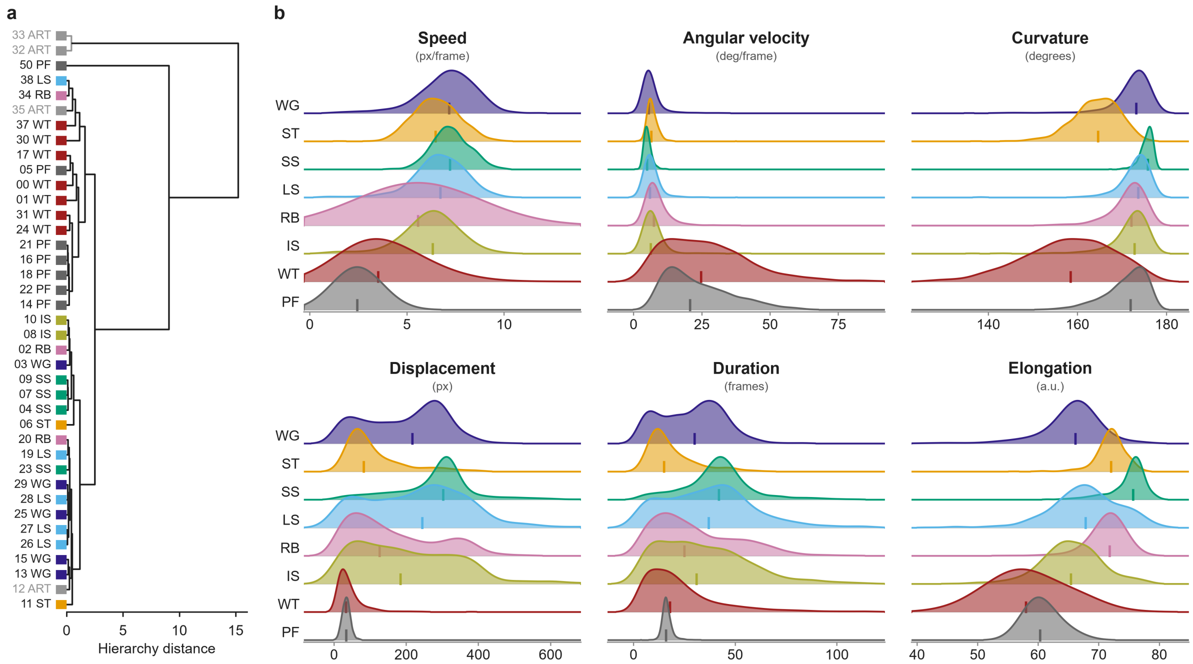


**Supplementary Figure 6:** Hierarchy and kinematic feature distributions of the Keypoint-MoSeq syllables. (a) Dendrogram of the hierarchical similarity between 39 syllable states, from which the 35 retained states were merged into the eight syllables; four artifact states (12, 32, 33 and 35, grey) belong to no syllable. Leaves in the order of the stored linkage, each labelled with its state ID, syllable abbreviation and colour; x axis, hierarchy distance. (b) Density of six kinematic features (speed, angular velocity, curvature, displacement, duration and elongation) for each of the eight syllables (13,437 bouts from 42 mice, LA n = 24, HA n = 18), one row per syllable in the order of Figure 4c; vertical tick, median. Abbreviations as in Figure 4c.

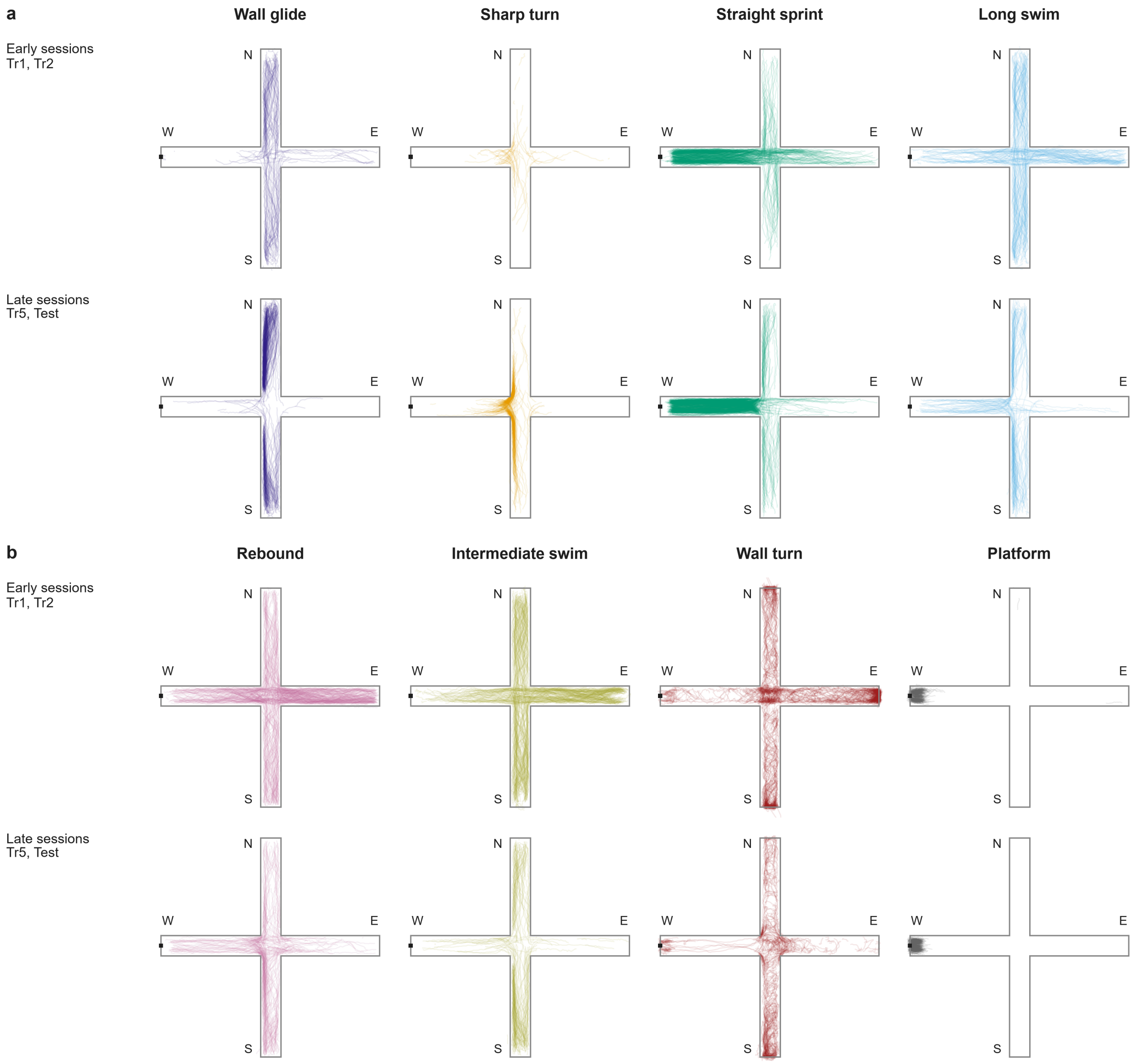


**Supplementary Figure 7:** Spatial deployment of each Keypoint-MoSeq syllable in early and late training. (a) Centroid paths of the bouts of Wall glide, Sharp turn, Straight sprint and Long swim on the Water Cross Maze, one maze per syllable, the early sessions (Tr1, Tr2; upper row) over the late sessions (Tr5, Test; lower row). (b) As (a) for Rebound, Intermediate swim, Wall turn and Platform. Each line is one bout (the bouts of Figure 5b,c, from 42 mice, LA n = 24, HA n = 18), coloured by syllable as in Figure 5; N, S, W, E, the maze arms; black square, the end of the west arm.

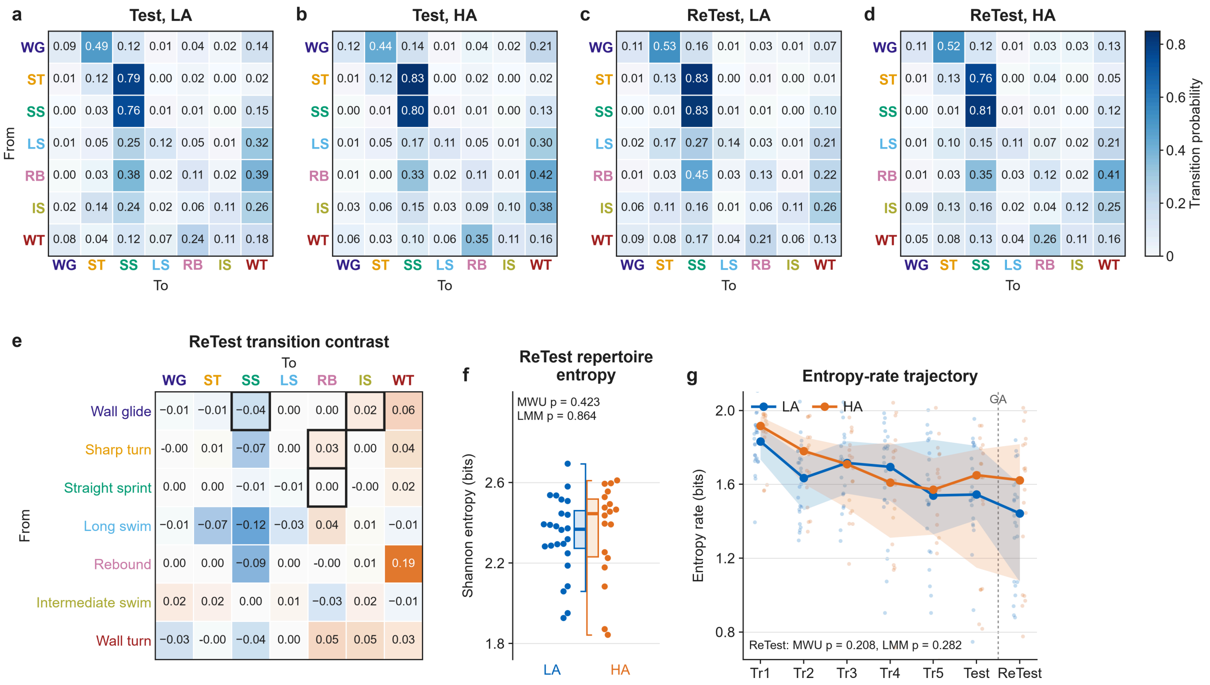


**Supplementary Figure 8:** Syllable transition matrices and entropy metrics in LA and HA mice. (a-d) Group median conditional transition probability matrices at Test for LA (a) and HA (b) and at ReTest for LA (c) and HA (d) for low anxiety (LA, blue, n = 24) and high anxiety (HA, orange, n = 18) mice; rows, current syllable; columns, next syllable; WG, Wall Glide; ST, Sharp Turn; SS, Straight Sprint; LS, Long Swim; RB, Rebound; IS, Intermediate Swim; WT, Wall Turn; values printed per cell on one colour scale. (e) Median transition probability at ReTest, HA minus LA; blue, larger in LA; orange, larger in HA; outlined cells, uncorrected two-sided Mann-Whitney U test $p < 0.05$. (f) Shannon entropy of the syllable frequency vector at ReTest, in bits; MWU, two-sided Mann-Whitney U test; LMM, linear mixed model with batch as random intercept. (g) Entropy rate, in bits, across the five training days (Tr1 to Tr5), Test and ReTest; dashed line, sevoflurane anaesthesia (GA) between Test and ReTest. Faint points, individual mice; filled circles joined by lines, group medians; shaded bands, interquartile range (25th-75th percentile) across mice; half-boxes, median and interquartile range, whiskers to the most extreme observation within 1.5 interquartile ranges.

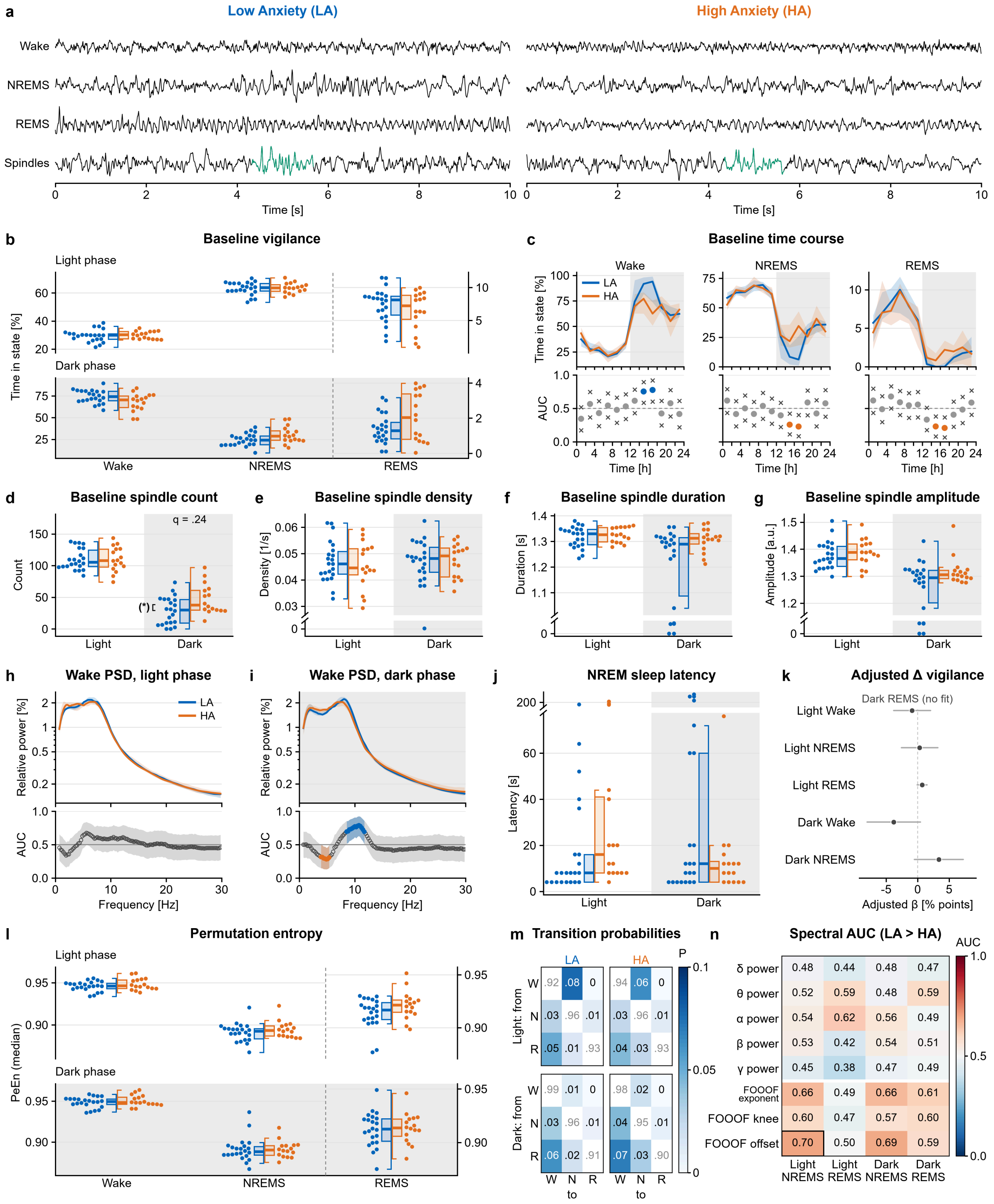


**Supplementary Figure 9:** Baseline sleep/wake architecture and spectral, latency, transition and complexity measures in LA and HA mice. (a) Representative 10-s EEG segments in WAKE, non-rapid eye movement sleep (NREMS) and rapid eye movement sleep (REMS) and a sleep spindle (green highlight), one LA and one HA mouse. (b) Percentage of time in WAKE, NREMS and REMS per phase of the baseline 24-h recording (bEEG; LA n = 22, HA n = 16). (c) Vigilance state percentages in 2-hour bins: line, group median; band, bootstrap 95% CI of the median; dark phase shaded. Lower strip, AUC (LA > HA), circles; 95% CI, crosses. Dots: grey at 0.3-0.7; otherwise blue (LA) or orange (HA) if CI excludes 0.5, black if not; dashed line, AUC = 0.5. (d-g) Sleep spindle metrics per phase: (d) count, (e) density, (f) median duration, (g) median normalized amplitude. (h, i) Relative power spectral density during WAKE per phase, 0.5-30 Hz: line, group median; band, 95% CI of the median. Lower strip, AUC (LA > HA) per frequency bin with 95% CI (grey band); bins whose CI excludes 0.5 filled in the higher-power group's colour. (j) Sleep latency (median of the first 5 inter-bout gaps) per phase. (k) Adjusted group effect (β, percentage points) with 95% CI from linear mixed models of the within-subject change in state percentages (Δ ~ group + baseline + (1 | batch)) per phase and state; dashed line, β = 0; dark REMS, no CI (random-effects term at zero). (l) Permutation entropy (median per mouse) per vigilance state and phase. (m) Markov transition matrices between states per phase and group; self-transitions in grey. (n) AUC (LA > HA) for power in five bands and for the fitting oscillations and one over f (FOOOF) aperiodic exponent, knee and offset during NREMS and REMS per phase; black frame, uncorrected $p < 0.05$. (b, d-g, j, l) Dots, individual mice; half-boxes, median and interquartile range, whiskers to the most extreme observation within 1.5 interquartile ranges. q, Benjamini-Hochberg FDR-corrected p of the two-sided Mann-Whitney U test; bracket with (*), uncorrected $p < 0.05$ only.

**Supplementary Table 1.** Group comparisons reported in the Results. Each row gives the measure, the number of mice contributing to that comparison, the effect size with its bootstrap confidence interval, and the significance values. Values in the prose are the medians and units that state each finding; the full inferential statistics are collected here. In the figure column a main-figure panel is written as 3c and a supplementary-figure panel as SF3c. This first part covers intra-anaesthetic EEG, conventional WCM performance, and syllable repertoire and sequence.

| Measure | n (LA / HA) | AUC (LA > HA) [95% CI] | p | Figure |
|---|---|---|---|---|
| **Intra-anaesthetic EEG** | | | | |
| Sevoflurane concentration at LORR and RORR, phase durations, burst-suppression architecture | 22 / 12 | range 0.34 to 0.64 | all MWU > 0.11 | |
| Band power, every band and maintenance phase | 22 / 12 | | none significant | |
| Absolute theta power, Adequate | 22 / 12 | 0.70 [0.52; 0.87] | MWU 0.054 | |
| Relative beta change across LORR | 22 / 12 | 0.23 [0.08; 0.41] | MWU 0.010; LMM < 0.001; batch ICC 0.21 | SF3c |
| Relative delta change across LORR | 22 / 12 | 0.74 [0.56; 0.90] | LMM 0.024 | SF3c |
| Relative gamma change across LORR | 22 / 12 | 0.30 [0.12; 0.50] | LMM 0.026 | SF3c |
| Aperiodic spectral edge frequency (SEF95), Adequate | 22 / 12 | 0.15 [0.05; 0.30] | MWU 0.001; FDR q 0.012; LMM 0.006 | 3b, 3c |
| Aperiodic spectral entropy, Adequate | 22 / 12 | 0.19 [0.06; 0.36] | LMM 0.010 | 3d |
| Aperiodic beta ratio, Adequate | 22 / 12 | 0.22 [0.08; 0.39] | LMM 0.014 | 3e |
| Aperiodic exponent, Adequate | 22 / 12 | | not significant | 3f |
| Aperiodic offset, Adequate | 22 / 12 | | not significant | 3g |
| Permutation entropy (PeEn), Adequate | 22 / 12 | 0.26 [0.10; 0.44] | MWU 0.022; LMM 0.051; batch ICC 0.60 | SF3e |
| Lempel-Ziv complexity (LZC), Adequate | 22 / 12 | 0.23 [0.08; 0.41] | | SF3e |
| Peak bandwidth, Discontinuity | 18 / 11 | 0.90 [0.75; 1.00] | MWU < 0.001; FDR q 0.012 | 3h |
| SEF95, pre-RORR window from 71 s to 47 s before RORR | 19 / 8 | | cluster permutation 0.031 | 3j |
| PeEn, pre-RORR and post-RORR windows | 19 / 8 | | cluster permutation, not significant (pre-RORR 0.59, post-RORR 0.18) | SF3d |
| **Conventional WCM performance** | | | | |
| Escape latency, Tr1 | 24 / 18 | 0.28 [0.14; 0.45] | MWU 0.019 | SF4a |
| Escape latency, Tr5 | 24 / 18 | | MWU 0.147 | SF4a |
| Wrong-platform visits, pre-anaesthesia Test | 24 / 18 | | MWU 0.018; FDR q 0.23 | SF4g, SF4j |
| Escape latency, ReTest | 24 / 18 | 0.30 [0.15; 0.48] | MWU 0.033; LMM 0.009; batch ICC 0.17 | SF4b, SF4j |
| Choice accuracy, ReTest | 24 / 18 | | not significant | SF4d, SF4j |
| Wrong-platform visits and start-arm bias, ReTest | 24 / 18 | | MWU 0.105, 0.169; none significant | SF4f, SF4h, SF4j |
| **Syllable repertoire and sequence** | | | | |
| Shannon entropy of the syllable frequency vector, every session | 24 / 18 | | all MWU > 0.36 | SF8f |
| Whole-matrix transition distance, ReTest | 24 / 18 | 0.62 [0.41; 0.80] | MWU 0.199 | |
| Wall Glide to Straight Sprint, ReTest | 24 / 18 | 0.72 [0.56; 0.86] | MWU 0.015; LMM 0.022 | 6e |
| Sharp Turn to Rebound, ReTest | 24 / 18 | 0.30 [0.15; 0.47] | MWU 0.028; LMM 0.027 | 6g |
| Straight Sprint to Rebound, ReTest | 24 / 18 | 0.29 [0.15; 0.46] | MWU 0.024; LMM 0.055 | 6h |
| Wall Glide to Intermediate Swim, ReTest | 24 / 18 | 0.30 [0.13; 0.49] | MWU 0.026; LMM 0.006 | 6f |
| Per-cell correction across the 7 x 7 matrix | 24 / 18 | | 0 of 49 significant at ReTest, 0 of 49 at Test | |
| Within-subject change, Test to ReTest, three transition cells | 24 / 18 | | 0.041, 0.008, 0.006 | |
| Within-subject change, Wall Glide to Intermediate Swim | 24 / 18 | | 0.292 | |
| Wall Turn bout duration, ReTest | 24 / 16 | 0.28 [0.12; 0.46] | MWU 0.020; LMM 0.038 | 6i |
| Wall Turn bout path length, ReTest | 24 / 16 | 0.24 [0.09; 0.41] | MWU 0.007; LMM 0.231 | 6j |
| Wall Turn swim speed, ReTest | 24 / 16 | | MWU 0.384; LMM 0.383 | |
| Wall Turn frequency, ReTest | 24 / 18 | | MWU 0.151; LMM 0.081 | |

**Supplementary Table 1, continued.** Syllable transitions from Tr5 to ReTest. The four transitions that differed between LA and HA mice at ReTest, compared at Tr5 and at Test, as the within-mouse change from Tr5 to Test, and as the difference between the change from Test to ReTest and the change from Tr5 to Test. Columns as in the first part.

| Measure | n (LA / HA) | AUC (LA > HA) [95% CI] | p | Figure |
|---|---|---|---|---|
| **Syllable transitions from Tr5 to ReTest** | | | | |
| Wall Glide to Straight Sprint, Tr5 | 24 / 18 | 0.56 [0.38; 0.73] | MWU 0.517; FDR q 0.99; LMM 0.773 | |
| Sharp Turn to Rebound, Tr5 | 24 / 18 | 0.46 [0.28; 0.64] | MWU 0.638; FDR q 0.99; LMM 0.412 | |
| Straight Sprint to Rebound, Tr5 | 24 / 18 | 0.45 [0.27; 0.63] | MWU 0.602; FDR q 0.99; LMM 0.390 | |
| Wall Glide to Intermediate Swim, Tr5 | 24 / 18 | 0.47 [0.28; 0.66] | MWU 0.732; FDR q 0.99; LMM 0.180 | |
| Wall Glide to Straight Sprint, Test | 24 / 18 | 0.43 [0.25; 0.61] | MWU 0.438; FDR q 0.95; LMM 0.857 | |
| Sharp Turn to Rebound, Test | 24 / 18 | 0.54 [0.36; 0.72] | MWU 0.656; FDR q 0.95; LMM 0.523 | |
| Straight Sprint to Rebound, Test | 24 / 18 | 0.56 [0.38; 0.73] | MWU 0.501; FDR q 0.95; LMM 0.170 | |
| Wall Glide to Intermediate Swim, Test | 24 / 18 | 0.44 [0.26; 0.63] | MWU 0.485; FDR q 0.95; LMM 0.329 | |
| Wall Glide to Straight Sprint, change Tr5 to Test | 24 / 18 | 0.39 [0.22; 0.58] | MWU 0.247; FDR q 0.73; LMM 0.923 | |
| Sharp Turn to Rebound, change Tr5 to Test | 24 / 18 | 0.60 [0.42; 0.77] | MWU 0.258; FDR q 0.73; LMM 0.187 | |
| Straight Sprint to Rebound, change Tr5 to Test | 24 / 18 | 0.57 [0.39; 0.75] | MWU 0.453; FDR q 0.94; LMM 0.687 | |
| Wall Glide to Intermediate Swim, change Tr5 to Test | 24 / 18 | 0.53 [0.35; 0.72] | MWU 0.712; FDR q 0.99; LMM 0.245 | |
| Wall Glide to Straight Sprint, change Test to ReTest minus change Tr5 to Test | 24 / 18 | 0.67 [0.49; 0.83] | MWU 0.069; FDR q 0.56; LMM 0.325 | |
| Sharp Turn to Rebound, change Test to ReTest minus change Tr5 to Test | 24 / 18 | 0.29 [0.15; 0.47] | MWU 0.024; FDR q 0.26; LMM 0.133 | |
| Straight Sprint to Rebound, change Test to ReTest minus change Tr5 to Test | 24 / 18 | 0.30 [0.14; 0.47] | MWU 0.026; FDR q 0.26; LMM 0.047 | |
| Wall Glide to Intermediate Swim, change Test to ReTest minus change Tr5 to Test | 24 / 18 | 0.38 [0.21; 0.57] | MWU 0.208; FDR q 0.89; LMM 0.040 | |

**Supplementary Table 1, continued.** Baseline sleep and post-anaesthetic change from each animal’s own baseline. Columns as in the first part; abbreviations are defined below the table.

| Measure | n (LA / HA) | AUC (LA > HA) [95% CI] | p | Figure |
|---|---|---|---|---|
| **Baseline sleep** | | | | |
| Dark-phase REMS proportion | 22 / 16 | 0.38 [0.18; 0.58] | MWU 0.204; LMM 0.026 | SF9b, SF9c |
| Dark-phase NREMS proportion | 22 / 16 | | MWU 0.084; LMM 0.054 | |
| Dark-phase WAKE proportion | 22 / 16 | | MWU 0.074; LMM 0.049 | |
| Light-phase proportions, all three states | 22 / 16 | | none significant | |
| NREMS latency at light onset | 22 / 16 | 0.31 [0.16; 0.49] | MWU 0.050; LMM 0.200 | SF9j |
| Dark-phase WAKE alpha power | 22 / 16 | 0.74 [0.57; 0.88] | MWU 0.012; LMM 0.024 | SF9h, SF9i |
| Light-phase WAKE theta power | 22 / 16 | 0.68 [0.50; 0.85] | MWU 0.060; LMM 0.208 | SF9h, SF9i |
| NREMS and REMS spectra, all five bands | 22 / 16 | | none significant | |
| Aperiodic offset, light-phase NREMS | 22 / 16 | | MWU 0.043; LMM 0.259 | SF9n |
| Dark-phase spindle count per 12 h | 22 / 16 | 0.30 [0.14; 0.48] | MWU 0.036; LMM 0.023 | SF9d |
| Spindle density, duration and amplitude | 22 / 16 | | none significant | SF9e to SF9g |
| Vigilance-state transitions, 12 comparisons | 22 / 16 | | all FDR q > 0.50 | SF9m |
| **Post-anaesthetic change from each animal's own baseline** | | | | |
| Dark-phase REMS change | 19 / 12 | 0.82 [0.65; 0.94] | LMM 0.007; baseline-adjusted model, no p (random-effects term at zero) | 7g, 7h |
| Dark-phase NREMS change | 19 / 12 | 0.75 [0.54; 0.92] | MWU 0.024; LMM 0.007; baseline-adjusted LMM 0.091 | 7g, 7h |
| Dark-phase WAKE change | 19 / 12 | 0.23 [0.07; 0.43] | MWU 0.014; LMM 0.004; baseline-adjusted LMM 0.078 | 7g, 7h |
| Light-phase REMS change | 19 / 12 | | MWU 0.25; LMM 0.049; baseline-adjusted LMM 0.072 | |
| Light-phase WAKE change | 19 / 12 | | MWU 0.45; LMM 0.044; baseline-adjusted LMM 0.552 | |
| Dark-phase spindle count change | 19 / 12 | 0.73 [0.52; 0.90] | LMM 0.022 | 7i |
| Spindle density, duration and amplitude change | 19 / 12 | | none significant | 7j to 7l |

AUC is the area under the receiver operating characteristic curve with its bootstrap 95% confidence interval, reported throughout in the low-anxiety greater than high-anxiety orientation, so a value above 0.5 means the low-anxiety group was larger and a value below 0.5 means the high-anxiety group was larger. MWU is the Mann-Whitney U test, LMM a linear mixed model with batch as a random intercept, ICC the batch intraclass correlation from that model, and FDR q a p value corrected for the false discovery rate. Baseline-adjusted models add each animal's own baseline value as a covariate. Empty cells mark comparisons for which that quantity was not computed. LA, low anxiety; HA, high anxiety; LORR, loss of righting reflex; RORR, recovery of righting reflex.
